# Strategies for Mitigating Impacts to Birds and Bats from Offshore Wind Energy Development: Available Evidence and Data Gaps

**DOI:** 10.1101/2024.08.20.608845

**Authors:** Julia Gulka, Steve Knapp, Anna Soccorsi, Stephanie Avery-Gomm, Paul Knaga, Kathryn A. Williams

## Abstract

A diversity of approaches exist for mitigating (e.g., avoiding, minimizing, or compensating for) the effects of offshore wind energy (OSW) development on birds and bats, but little is known about the effectiveness of many of these approaches. To address this knowledge gap, we reviewed the scientific and gray literature to evaluate the evidence base for potential bird- and bat-related mitigation approaches for OSW, including studies from other industries where relevant (e.g., terrestrial wind energy, offshore oil and gas industry). Of a total of 219 mitigation approaches, most focused on minimization, with far fewer addressing avoidance or compensation. Sixty-five percent had no evidence of testing. Of the 76 mitigation approaches that were field tested or implemented, we found evidence of effectiveness for only 40 approaches, of which only 10 were specific to the OSW sector. Consequently, a majority of all the mitigation approaches for birds (82%) and bats (89%) lacked any evidence of effectiveness. For birds, minimization approaches related to lighting reduction were the most tested and effective methods for reducing maladaptive attraction and collisions. For bats, minimization approaches involving adjustments to turbine operations (e.g., curtailment of turbine blades) were most tested and effective methods for reducing collisions. Given the limited evidence of effectiveness for most mitigation approaches, assuming their success systematically underestimating project-level and cumulative impacts, underscoring the need to prioritize avoidance, consistent with the mitigation hierarchy, and rigorously test mitigation approaches for birds and bats in offshore environments.

**RÉSUMÉ:** Il existe une diversité d’approches pour atténuer (e.g. éviter, minimiser ou compenser) les effets du développement de l’ l’Énergie Éolienne Extracôtière (EEE) sur les oiseaux et les chauves-souris, mais l’efficacité de nombreuses de ces approches est encore peu connue. Pour combler ces lacunes, nous avons passé en revue la littérature scientifique et la littérature grise afin d’évaluer les données probantes concernant les mesures d’atténuation potentielle pour l’EEE et les oiseaux et les chauves-souris,, inclus des études d’autres industries lorsque cela était pertinent (e.g., l’énergie éolienne terrestre, l’industrie pétrolière et gazière extracôtière). Sur un total de 219 approches d’atténuation pertinentes pour l’EEE, la plupart des approches se concentraient sur la minimisation, avec beaucoup moins d’approches portant sur l’évitement ou la compensation. Soixante-et-un pourcent des approches proposées ne semblent pas avoir été testées. Parmi les 76 approches d’atténuation qui ont été testées sur le terrain ou mises en œuvre, nous avons trouvé des preuves de leur efficacité pour seulement 40 approches, dont seulement 10 étaient spécifiques àle secteur de l’EEE. Par conséquent, la majorité des approches d’atténuation pour les oiseaux (82 %) et les chauves-souris (89%) ne disposaient d’aucune preuve d’efficacité. Chez les oiseaux, les approches de minimisation liées à la réduction de l’éclairage étaient les méthodes les plus couramment testées et les plus efficaces pour réduire l’attraction et les collisions. Chez les chauves-souris, les approches de minimisation impliquant la modification du fonctionnement des turbines (e.g., réduction de la vitesse de rotation ou mise en drapeau des hélices) étaient les méthodes les plus couramment testées et les plus efficaces pour réduire les collisions. Compte tenu des preuves limitées de l’efficacité de la plupart des approches d’atténuation, le fait de présumer de leur succès conduit à une sous-estimation systématique des impacts à l’échelle des projets et des impacts cumulatifs, ce qui souligne la nécessité de privilégier l’évitement, conformément à la hiérarchie d’atténuation, et de tester rigoureusement les mesures d’atténuation pour les oiseaux et les chauves-souris en milieux offshore.

**IMPLICATIONS FOR MANAGERS:**

- Based on our literature review of potential mitigation approaches for birds and bats in relation to offshore wind development, most approaches remain untested or lack evidence of effectiveness (bats – 89%, birds – 82%).
- Most minimization approaches have primarily been tested in the terrestrial context, leaving important questions regarding the transferability of these results to the offshore environment.
- Given the limited evidence of effectiveness for most mitigation approaches, assuming their success systematically underestimates project-level and cumulative impacts, underscoring the need to prioritize avoidance and rigorously test mitigation approaches for birds and bats in offshore environments.

## INTRODUCTION

Offshore wind energy (OSW) development is a vital strategy for reducing global reliance on fossil fuels but also poses environmental challenges that require mitigation (Peste et al. 2015). Many bird and bat species worldwide are in decline and protected at state, provincial, federal, or international levels (Dias et al. 2019, Frick et al. 2020, Neate-Clegg et al. 2020). Within this context, it is critical to consider the potential effects of OSW developments on these wildlife taxa. There are three main types of effects to birds and bats from OSW development: collisions, behavior change (e.g., avoidance, attraction), and habitat-mediated change (e.g., changes in prey; Williams et al. 2024). Although our scientific understanding of the scale of these effects varies widely and significant gaps remain, several comprehensive reviews have synthesized the current state of knowledge on how OSW development affects birds and bats (Marques et al. 2014, Goodale and Stenhouse 2016, Fox and Petersen 2019, Solick and Newman 2021, Lamb et al. 2024, Williams et al. 2024). These reviews highlight pathways of effects that can inform mitigation strategies.

Currently, a variety of mitigation approaches are proposed and used by the OSW industry, in many cases drawing from other industries (e.g., terrestrial wind energy, other maritime industries), but there is limited information about the effectiveness of many approaches to mitigate the effects of developments on birds and bats. Given the harsh and inaccessible nature of the offshore environment, it can be both challenging and expensive to test mitigation strategies. Moreover, high environmental variability, often coupled with low sample sizes for studies that are trying to measure change in these systems, create challenges in achieving adequate statistical power to provide evidence of effectiveness (Maclean et al. 2013, Regional Synthesis Workgroup of the Environmental Technical Working Group 2023). Additionally, wildlife species and their habitats may be affected by a range of stressors, and the sensitivity and vulnerability of species varies in relation to a variety of factors, both internal (e.g., demographics, life history), and external (e.g., facility layout, location; Lamb et al. 2024, Williams et al. 2024), further complicating testing of mitigation approaches in situ. Consequently, some mitigation approaches may be implemented without sufficient evidence of their effectiveness, potentially wasting resources and creating a false sense of confidence in their ability to reduce the effects of OSW development.

### Overview of offshore wind effects to birds and bats

Implementing effective mitigation requires an understanding of the stressors to wildlife, potential responses, and mechanisms driving those responses. Key effects associated with offshore wind developments are those caused by the presence of permanent infrastructures, including collisions of birds and bats with turbine blades, changes in behavior (e.g., attraction to or avoidance of wind farms), and habitat-mediated changes (e.g., increased prey near turbines). Collisions of migrating birds (e.g., songbirds, raptors) and bats are well documented at onshore facilities (Allison et al. 2019) and concerns over high mortality levels, particularly for migratory bats, has driven significant efforts to monitor and mitigate collision risks in the terrestrial wind industry (Adams et al. 2021, Whitby et al. 2024). Offshore, monitoring collisions is far more challenging; carcass searches, a key detection method onshore, are largely impossible in the marine environment. Existing evidence, while limited, suggests that collisions of marine birds with OSW facilities in Europe (documented via direct observation and camera systems) appear to be much rarer than in terrestrial contexts (Skov et al. 2018, Tjørnløv et al. 2023), though the extent of this phenomenon remains poorly documented.

Behavioral responses can include avoidance at multiple scales and/or attraction of animals due to the presence of structures and associated vessels (Williams et al. 2024). Behavioral change has been documented for a range of marine bird species at OSW facilities in Europe, with the type and degree of attraction or avoidance depending on many factors including species, facility location, and design (Dierschke et al. 2016, Lamb et al. 2024). Attraction may be due to increased opportunities for perching and roosting (Dierschke et al. 2016), lighting on turbines and substations (Rebke et al. 2019), or habitat-mediated change (e.g., increased prey; Russell et al. 2014). The fitness consequences of attraction to wind facilities are unclear; while there is the potential for positive energetic consequences, such attraction is also thought to increase collision risk. Some species of birds may be attracted to and disoriented by artificial light at night, particularly during migration; this can lead to death via collisions or exhaustion (Montevecchi 2006, Burt et al. 2023). For bats, artificial lighting could be an attractant, in part due to the attraction of insects on which bats feed (Stone et al. 2015, Jonasson et al. 2024), though a recent review suggests that this mechanism is likely not a major source of bat attraction to wind turbines (Guest et al. 2022), with lighting potentially causing avoidance for some but not other bat species.

Avoidance of individual turbines within the wind farm footprint or the entire offshore wind facility with resultant changes in distributions (e.g., displacement) may relate to the presence of the offshore wind facility infrastructure itself, associated activities, such as vessel traffic (Cook et al. 2018), or artificial light (Syposz et al. 2021). Avoidance presumably decreases any risk of collision; while it is thought to also have potential negative energetic consequences, evidence for fitness consequences is currently limited (Garthe et al. 2023). Another way OSW development may affect birds and bats is via changes in the physical environment and/or resource availability (i.e., habitat-mediated effects; Williams et al. 2024), which can include changes in oceanographic conditions and prey (e.g., artificial reef effect; Degraer et al., 2020). These altered conditions may change the distribution, abundance, and structure of prey communities.

### The mitigation hierarchy

Mitigation of effects such as those posed to birds and bats by the OSW industry is guided by the well-established mitigation hierarchy, which prioritizes measures that avoid potential effects altogether as the most effective strategy, followed by minimization of unavoidable effects, and finally, compensation for residual effects that cannot be avoided or minimized (Council on Environmental Quality 2020).

This hierarchy provides a structured framework for addressing the effects of anthropogenic activities, ensuring that efforts first aim to prevent harm before attempting to reduce or compensate for it. Hereafter, the term ‘mitigation’ is used to encompass all three steps of this hierarchy.

Avoidance measures are generally implemented during early planning phases of OSW development and typically involve siting development activity away from regions with high risks (Winiarski et al. 2014, Croll et al. 2022). Minimization primarily occurs during pre-construction and operational phases and may include adjustments to facility components or operational parameters.

Compensation, the final step, involves activities designed to offset effects that cannot be avoided or minimized, and may include approaches such as creating or improving habitat (Wolf et al. 2006, Croll et al. 2022, Spatz et al. 2022).

### Study Objectives

Existing efforts have examined the effectiveness of various mitigation approaches, a key example being the Conservation Evidence project (https://www.conservationevidence.com/). While not offshore wind-specific, this international project takes a science-based approach to assessing a wide range of biodiversity conservation measures and includes links to the scientific studies where mitigation effectiveness was examined. Several other efforts have focused specifically on assessing the effectiveness of mitigation approaches for OSW development, though these have typically not focused on birds and bats (for example, see Verfuss et al., 2016). One recent assessment to inform OSW development in Scotland (Crown Estate Scotland 2024) included birds but was unable to find data on the effectiveness of mitigation approaches in most cases and thus chose to define mitigation success as situations in which “no deleterious effects were recorded”.

The goal of this review is to fill this gap in the scientific literature and focus explicitly on assessing the state of knowledge regarding the effectiveness of mitigation for birds and bats in relation to OSW development. We compile an inventory of mitigation approaches that have been proposed and are potentially relevant for use for OSW development across jurisdictions. Because many OSW mitigation approaches have been sourced from related industries, most notably the terrestrial wind industry, these industries are included in this assessment, though our focus is on mitigation for aspects of OSW developments in the marine environment (e.g., project footprint, cable and transit routes) as opposed to onshore aspects (e.g., transmission interconnection). In this study we summarize the weight of evidence supporting various mitigation approaches, identify key gaps in our understanding, and use this information to prioritize future research to further examine the effectiveness of mitigation options for birds and bats in relation to OSW development.

## METHODS

To assess the state of knowledge regarding the testing and demonstrated effectiveness of mitigation approaches for birds and bats associated with OSW development, we compiled and evaluated mitigation approaches described in the peer-reviewed and gray literature, which constitute the primary evidence base used in OSW regulatory decision making (Szostek et al. 2024). Using the Mitigation Practices Database Tool and an updated literature review through 2023 (described below), we synthesized mitigation approaches into discrete categories and classified each approach based on whether it had been tested or implemented and whether evidence supported its effectiveness. This study does not quantify the magnitude of impact reduction achieved by individual mitigation approaches; rather, it evaluates whether mitigation approaches have been tested and whether evidence exists to demonstrate effectiveness.

### Mitigation Practices Database

In 2018-2019, the New York State Energy Research and Development Authority funded a desktop review to aggregate and assess practices for mitigation of effects to fisheries and offshore species (i.e., birds, bats, marine mammals, sea turtles, fish, and benthos) associated with all phases of OSW development. At the time of this review, this database, the Mitigation Practices Database (MPD) Tool (https://www.nyetwg.com/mpd-tool), included sources (peer-reviewed and gray literature) relevant to birds and bats (n=107) identified via Google Scholar (search terms: offshore wind + birds + mitigation; offshore wind + bats + mitigation) and via an internal Biodiversity Research Institute Mendeley database (search terms: mitigation; mitigate; minimize). The structure of the database included information on the taxonomic group for which the mitigation was proposed or tested, the types of stressors and effects that were addressed, the phase of development, and evidence of effectiveness for the mitigation approach, among other information (Table 1).

**Table 1.** Key types of information collated in review of mitigation options available for offshore wind development for birds and bats, synthesized from the existing published and gray literature.

| Type of Information | Options | Definition |
| --- | --- | --- |
| Mitigation Category | Siting/Activity Restriction, Turbine Operation Parameters, Other Operations Parameters, Structure Configuration, Deterrence and Attraction Reduction, Lighting and Visibility, Compensation | High level categorization of mitigation approach |
| Resource | All Birds, All Bats, Marine Birds, Nocturnal Aerial Migrants | Functional species group for which mitigation approach is relevant, as defined in source document(s) |
| Stressors | Bottom Disturbance, Changes in Vessel Traffic, Light, Long-term Structures, Sound, Vibration, Water Quality Changes | External stimuli that can cause changes to the behavioral, physical, chemical, and/or biological characteristics of an organism or species, or the ecosystem inhabited by the organism/species |
| Potential Effects | Behavioral Disturbance, Displacement, Attraction, Habitat Fragmentation/Modification, Injury/Mortality, Community Alteration/Invasive Species | The changes to the behavioral, physical, chemical, and/or biological characteristics of an organism or species, or the ecosystem inhabited by the organism/species, due to stressors related to OSW development |
| Development Phase | Pre-construction, Construction, Operations and Maintenance, Decommissioning | Stage of offshore wind facility development/operation for which the approach is relevant, each of which encompass a number of activities and, as a result, may have different types of stressors |
| Industry Origin | Offshore Wind, Terrestrial Wind, Oil and Gas, Maritime, Generic/Other | Type of industry for which mitigation practice was suggested or implemented. Generic/Other used if unspecified or another industry (e.g., agriculture) |
| Mitigation Hierarchy | Avoidance, Minimization, Offset | Level of the mitigation hierarchy |
| Implementation Status | Not Implemented, Field Tested, Implemented, Implemented and Evidence of Effectiveness, Unknown | The degree to which the use or effectiveness of a mitigation practice was tested in the cited literature |

### Update to literature review

We conducted an additional, more comprehensive literature review in early 2024 to add sources not identified in the original database and to update information on mitigation approaches for birds and bats from 2019-2023. A series of literature searches were conducted using Google Scholar (search terms: bird+wind+mitigation; bat+wind+mitigat*; bat+wind+mitigat*+offshore; bird*+wind+mitigat*+offshore; bird+wind+compensation; bat+wind+compensation; bird+wind+offsets; bat+wind+offsets; bird+offshore wind+compensation; bat+offshore wind+compensation; bird+offshore wind+offsets; bat+offshore wind+offsets; bird+wind+avoidance; bat+wind+avoidance; bird+wind+siting; bat+wind+siting), the Tethys Knowledge Base (Filters: Wind energy content + birds or bats; search terms “mitigation”, “compensation”, “avoidance”), and the Renewable Energy Wildlife Institute Research Hub (Filters: Offshore wind and wildlife; Bat research; Raptor research; search terms: “mitigation”, “compensation”, “avoidance”. These search methods were chosen to replicate the MPD Tool review process as best we could while also ensuring that the search was as comprehensive as possible and key research texts from both the gray and published literature were included.

Following identification of sources, the abstract or summary of each source was manually reviewed for initial relevance. If there was no mention of mitigation, avoidance, minimization, compensation, or offsetting, sources were deemed not relevant and were not reviewed further. The remaining sources were reviewed, and information was manually extracted following a similar structure to the MPD Tool, above. This initial filtering resulted in the identification of 141 additional sources, of which 93 were included in the literature review (the remaining 48 were deemed not relevant following a more in-depth review of the entire document). Combined with approaches from the MPD Tool, there were a total of 201 sources (n=127 peer reviewed, n= 74 gray literature). These sources focused on various industries, including offshore wind (n=77), terrestrial wind (n=104), offshore oil and gas (n=15), other maritime industries (n=7), and generic or other industries (e.g., agriculture; n=39). These new sources have since been incorporated into an updated version of the MPD Tool. Mitigation approaches were synthesized in the new database based on topical commonalities. For example, if multiple sources discussed variations on the mitigation strategy ‘paint turbine blades,’ they were synthesized into a single mitigation measure with study-specific details noted as relevant.

### Quantifying Evidence of Effectiveness

We examined the level of evidence from each source and assigned an “implementation status” for each mitigation approach of: 1) Not Implemented/Unknown, 2) Field Tested, and 3) Implemented. “Field testing” was defined as a small-scale experimental design (in-situ or ex-situ), while “implemented” meant that the strategy was adopted in a real-world setting at the development project scale. As a mitigation strategy may have come from multiple sources, categories were assigned based on the most advanced status. We then ranked the level of evidence for the effectiveness of each mitigation approach using the following categories: 1) None, 2) Limited, 3) Mixed, 4) Strong. These categories were specified separately for birds and bats. Effectiveness, in this context, was defined as the degree to which a mitigation approach was successful in producing the desired result, relying solely on information provided in the source document. This ranking scheme has a level of subjectivity and assumes scientific rigor in data collection and analysis, which may have been variable across studies. Any mitigation strategies that had an implementation status of Not Implemented/Unknown received a “None” ranking, along with those that were field tested or implemented with evidence that the strategy *did not* produce the anticipated results. Strategies received a “Limited” ranking if there were 1-2 small scale studies or field tests showing evidence of effectiveness, often for a particular species or context. Mitigation strategies received a “Mixed” ranking if there were one or multiple studies or field tests that showed contradictory results in achieving similar goals. Finally, a “Strong” ranking was applied to strategies with multiple sources (or in some cases reviews/meta-analyses) showing that the strategy produced the anticipated results. Because the level of testing in the offshore wind context was lacking, we did not distinguish effectiveness by industry but rather described these nuances regarding effectiveness context in the results.

### Limitations of the Literature Review

Given how various mitigation-related terms are generally used in the literature, the results primarily included minimization measures, with much less representation of avoidance or compensation approaches. Despite the prevalence of the mitigation hierarchy in the scientific literature, the term “mitigation” is sometimes used synonymously with “minimization,” and that was evident in the types of studies that were identified during our review. While we made an effort to find additional avoidance and compensation-related sources via targeted search terms, the scope of this review largely excluded marine spatial planning exercises (e.g., macro-siting), stakeholder engagement activities, and permitting documents developed by governmental agencies where siting-focused and compensation-focused mitigation tended to be discussed. Some of these activities (e.g., stakeholder engagement by government entities) also may not have been particularly well represented in the source documents, despite our best efforts to include gray literature in addition to peer-reviewed studies.

## RESULTS

A broad range of mitigation approaches (n=219) were identified from the existing literature (n=201 sources; Appendix A-B), focusing on a variety of stressors, potential effects, and industries. Many approaches were suggested by multiple sources (mean=2.3, range=1-19). There were a wider variety of mitigation approaches focused on birds (n=133) than bats (n=31), with some focused on both (n=55). About 14% of mitigation approaches were specific to marine birds or nocturnal avian migrants, but most were relatively general. Minimization approaches were the most common (n=123), followed by avoidance (n=72), with the fewest compensation/offset-related approaches (n=24). Most mitigation approaches came at least in part from the terrestrial wind industry (67%), with about 44% focused on offshore/marine environments, including offshore oil and gas, offshore wind, and other maritime industries (Figure 1a); the remainder were “general” mitigation approaches that were either not suggested in relation to a specific type of industrial development, or for an industry not listed above (e.g., agricultural sector).

**Figure 1.**
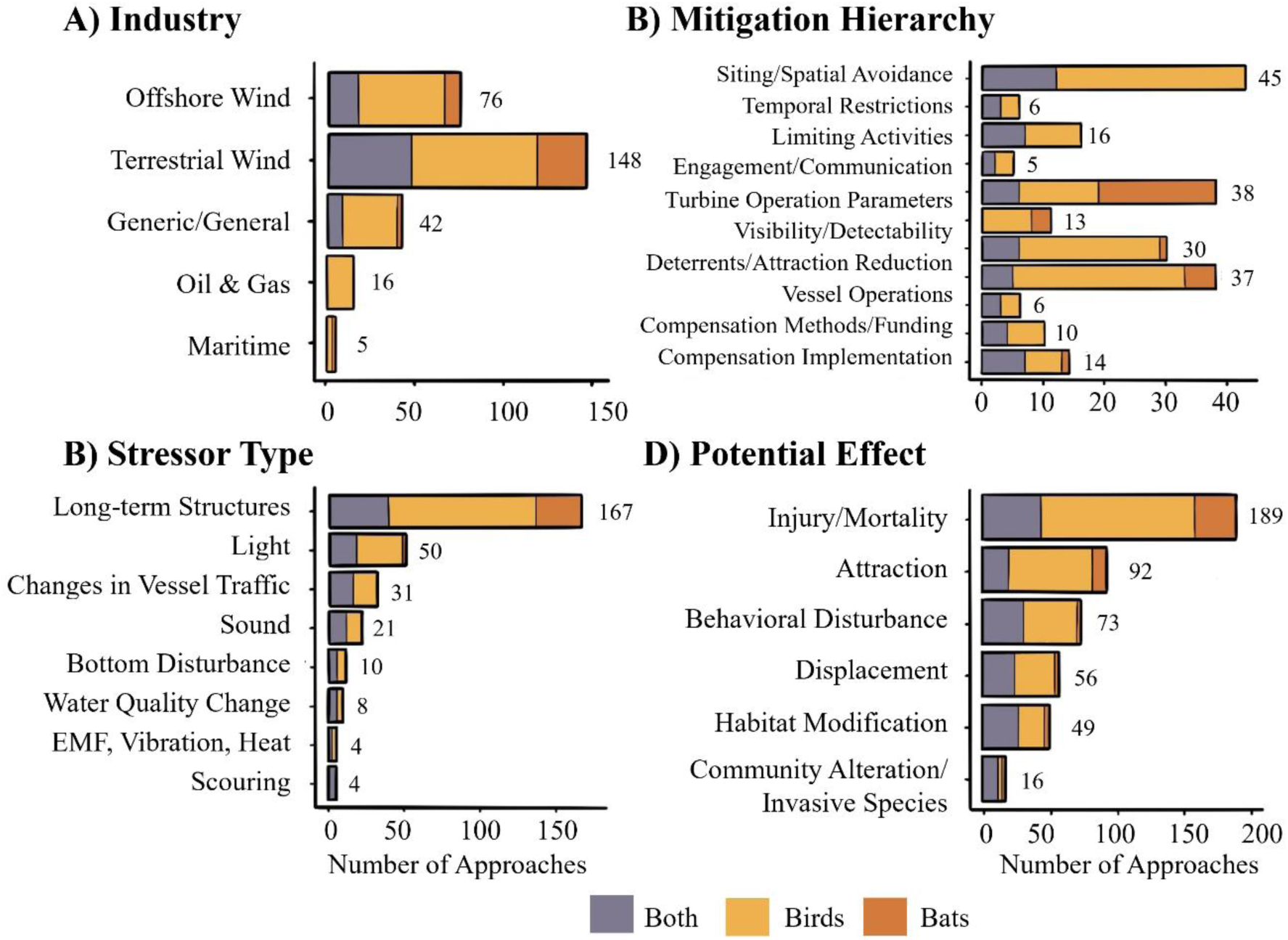
Number of bird and bat mitigation approaches by industry (A), mitigation category (B), stressor (C), and potential effect type (D). Mitigation approaches were directed towards birds (yellow), bats (orange), or both taxa (gray). Mitigation categories are mutually exclusive; however, each mitigation approach could be listed under multiple stressors, effects, and industries. Numbers indicate total approaches per category.

To aid in further summarization, we categorized mitigation approaches within the mitigation hierarchy (Figure 1b). Avoidance-related approaches were categorized into 1) siting and spatial avoidance (e.g., avoiding sensitive habitats, turbine configuration; n=45), 2) temporal restrictions (e.g., avoiding particular seasons/time periods, n=6), 3) limiting activities (e.g., avoiding pollutants, invasives introduction, etc.; n=16), and 4) engagement and communication (n=5). Minimization-related approaches were categorized into 1) turbine operation parameters (e.g., curtailment; n=38), 2) lighting to reduce attraction (e.g., changing color of lights; n=30), 3) visibility and detection enhancement (e.g., painting turbine blades; n=12), 4) deterrents and other attraction reduction measures (n=37), and 5) vessel operations (e.g., speed reductions; n=6). Finally, compensation-related approaches were categorized into 1) methods and funding (e.g., how to calculate offsets; n=10) and 2) compensation implementation (e.g., restoring similar habitats or species in other locations, reducing fisheries bycatch to offset collision mortality; n=14). The widest variety of suggested approaches were in the siting and spatial avoidance category (n=45), most of which were focused on birds (69%). The category of changes to turbine operations (n=38) included the greatest number of bat-related approaches (50%), mostly variations of curtailment, which is a key mitigation tool used for bats at terrestrial wind facilities (Adams et al. 2021, Whitby et al. 2024). Three quarters of mitigation approaches (76%) focused on the presence of long-term structures (e.g., turbines, substations) as a stressor type, followed by lighting (23%) and changes in vessel traffic (14%; Figure 1c). The bulk of mitigation approaches focused on mitigating injury/mortality (86%) or attraction (42%; Figure 1d). Within approaches focused on injury/mortality, almost half (46%) also focused on mitigating attraction given the potential links between these effects.

Surprisingly, only 76 (35%) mitigation approaches were implemented or field tested in at least one source document (Figure 2a). Of those 76 approaches, some level of evidence of effectiveness was found for 40 approaches for birds and 23 for bats - representing just 11-18% of all mitigation approaches identified in this review. This combines all levels of effectiveness (limited, mixed, strong), with only 5 approaches with strong evidence (2%; Figure 2b). Thus, 82-89% of mitigation approaches had no evidence of effectiveness based on the reviewed source documents, underscoring critical gaps in our understanding of appropriate mitigation. For the remaining 143 approaches (65%), it was clear that 26% had not been implemented or tested, while for the remaining 39% the implementation and testing status was unclear. Unsurprisingly, given the difficulties in testing mitigation approaches in the offshore environment, a slightly lower percentage of offshore-focused mitigation approaches (n=93, or about 33%) had been field tested or implemented as opposed to 39% of approaches targeted partially or wholly on terrestrial systems. Where mitigation approaches had been tested, it was typically at a small scale, which led to small sample sizes and other issues that may make it difficult to scale up results to understand the likely influence of mitigation approaches that are implemented more broadly.

**Figure 2.**
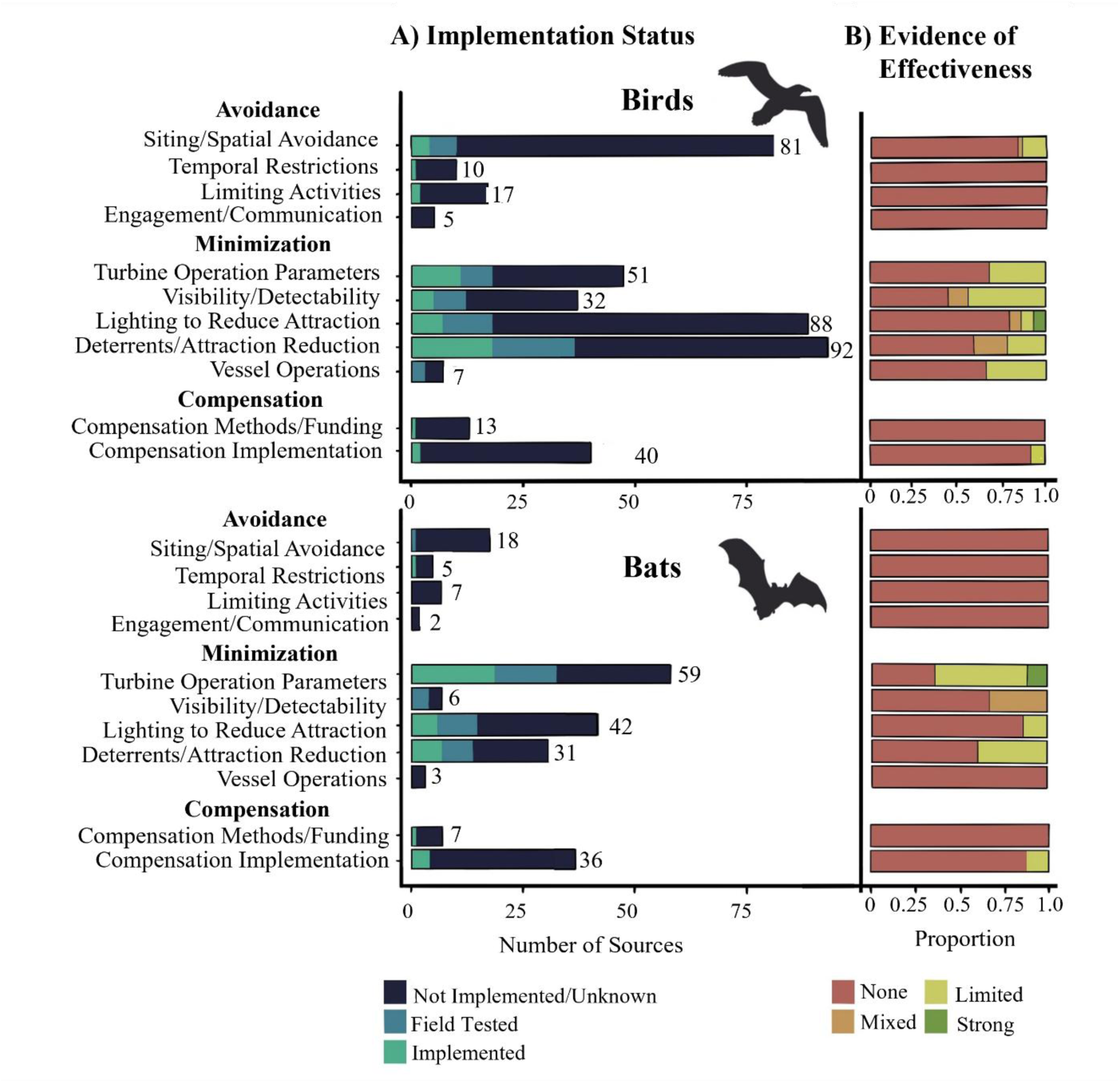
A) Number of sources by category for birds (top) and bats (bottom) for various mitigation approaches along with the level of testing and implementation (dark blue = mitigation approach was not implemented/tested in the source document or it was unclear whether it had been implemented; light blue = mitigation approach had been field tested in the source document, but not implemented in a real-world situation; green = mitigation approach was implemented in a real-world situation. Mitigation categories are mutually exclusive; however, if the same mitigation approach is discussed in multiple source documents within a category, it is represented multiple times in this figure (and may be classified into different effectiveness types, if the studies had differing results). Numbers indicate total number of sources per category. B) Proportional level of evidence of effectiveness for mitigation approaches by category. Effectiveness was categorized into 1) strong (green), 2) limited (yellow), mixed (orange) and none (red).

### Avoidance measures

#### Siting and spatial avoidance

The location and configuration of offshore wind facilities (and other anthropogenic infrastructure) have the potential to influence how birds and bats perceive these structures and therefore how they respond to and interact with them. Siting and other spatial-based avoidance approaches (n=45) included macro-siting (e.g., siting the entire wind farm away from important habitat use areas), micro-siting (e.g., placing individual turbines and other infrastructure within a project footprint in such a way as to avoid key use areas, and other spatial design components (Allison et al. 2019). Important habitat use areas were defined in the literature in a variety of ways, including known bat maternity roosts and hibernacula (American Wind Wildlife Institute 2019, Reusch et al. 2022), avian breeding colonies and nesting areas (May et al. 2015, Defingou et al. 2019, Simmons et al. 2020), designated Important Bird Areas and other areas of high bird usage, e.g., with known natural concentrations of birds (Langston and Pullan 2003, Drewitt and Langston 2006, Lüdeke 2019), known feeding, staging, or loafing locations (Defingou et al. 2019, Lüdeke 2019, Thaxter et al. 2019), migratory and commuting pathways (Langston and Pullan 2003, Mockrin and Gravenmier 2012, Defingou et al. 2019, Lüdeke 2019), areas of high prey density (Watson et al. 2018), shallow waters and waters in proximity to shore (Williams et al. 2015, Goodale and Milman 2020, Lagerveld et al. 2023), sensitive habitats (Mockrin and Gravenmier 2012, Rydell et al. 2012), and undisturbed areas (Mockrin and Gravenmier 2012, Eshleman and Elmore 2013). Several literature sources suggested using specific types of data to help define these important habitat use areas, such as bat acoustic activity data, environmental data, and weather radar data on bird and bat migration (Murgatroyd et al. 2021, Cohen et al. 2022, Gaultier et al. 2023). Micro-siting and design-based avoidance measures include adjusting the number of turbines, turbine layout (e.g., clustering, spacing), and turbine size (overall height, size of “air gap” below the lower edge of rotor-swept zone). Additional considerations related to clustering included leaving open migration corridors (Cook et al. 2011), leaving larger gaps between rows in the direction of prevailing winds (Tulp et al. 1999), and avoiding placing infrastructure within daily flight paths to foraging and breeding sites to ensure safe passage through or around the wind facility (Vaissière et al. 2014).

Few approaches in this category were tested for effectiveness. The only approaches for which there is limited evidence within this category (n=7) relate to turbine configuration and size (e.g., micro-siting and design) specifically for birds, with evidence primarily coming from the terrestrial wind context. Multiple studies found that increasing turbine height helped to reduce raptor mortalities at terrestrial wind facilities (Arnett and May 2016), including for Golden Eagles (*Aquila chrysaetos*; CEC 2002). However, the number of neotropical migrant mortalities was found in one study to be two orders of magnitude higher for 300m television towers compared with 90m towers (Crawford and Engstrom 2001). A recent study found rotor diameter was positively correlated with bird mortality but not with bat mortality rates at terrestrial wind farms, while decreasing ground clearance led to increased mortality rates for both birds and bats (Garvin et al. 2024). The effectiveness of altering turbine configuration varied by species and by the turbine characteristics that were altered, including overall height, distance between the surface of the water and the lower edge of the turbine blade (i.e., air gap), and rotor diameter (Garvin et al. 2024).

#### Temporal restrictions

In addition to spatial avoidance methods described above, several approaches focused on temporal avoidance (n=6), which involves scheduling disruptive activities to avoid sensitive life history periods and/or times of high habitat use. This included scheduling construction activities and maintenance activities (including vessel and helicopter activity) to avoid the avian breeding season and other times of high bird or bat abundance, such as migratory or wintering periods (Cook et al., 2011; English et al., 2017; Eshleman and Elmore, 2013; Gartman et al., 2016; Langston, 2013; May et al., 2015; Mockrin and Gravenmier, 2012; USFWS, 2021; Vaissière et al., 2014; World Bank Group, 2015). Seasonal limitations on construction activities to mitigate effects on marine birds have been implemented in multiple sources related to OSW development. For example, construction was limited to March-October at a European offshore wind farm to protect wintering birds, and restricted from December-March at another site to protect Common Scoter (*Melanitta nigra*; English et al. 2017). Despite this implementation, the effectiveness of such measures was not reported.

#### Limiting activities

Our review found additional avoidance methods focused on limiting activities (n=16), some of which (n=6) were targeted at marine birds specifically. These included sequencing and phasing turbine installation and other construction activities (English et al. 2017), limiting construction of artificial breeding structures in proximity to development (Everaert and Stienen 2007), co-locating construction equipment across projects (Drewitt and Langston 2008), and avoiding introduction of invasive species and pollution (USFWS 2012). Of the 16 strategies identified, only two were implemented, both targeting marine birds at OSW developments in the UK. In one case, turbine installation progressed from north to south over the course of a season to avoid an effect on a nearby Little Tern (*Sternula albifrons*) colony (English et al. 2017); in the other example, a phased and adaptive development approach was applied, with the second phase only permitted once the developer demonstrated no construction-related change to Red-throated Loon (*Gavia stellata*) habitat that would compromise identified conservation objectives (English et al. 2017). Effectiveness of these avoidance-based approaches was not reported.

#### Engagement and communication

While not exclusive to avoidance, approaches related to engagement and communication with stakeholders (n=5) were identified as potential mitigation strategies that were broadly applicable to both birds and bats. These approaches were included in the “avoidance” category because they typically related to early and standardized engagement with stakeholders and state and federal agencies, particularly in regard to siting, as a method of minimizing the potential for effects (USFWS, 2021). None of these strategies were tested for effectiveness within the reviewed literature, likely because the majority of these strategies were documented in guidance papers intended to provide planning assistance.

### Minimization measures

Minimization approaches were the most abundant (n=124; 56%) compared with avoidance and compensation and also had the highest levels of implementation and testing (Figure 2; Table 2).

**Table 2.** Minimization approaches for birds and bats relevant to offshore wind development and the level of evidence of effectiveness (none, limited, mixed, strong)

| Category | Goal | Subcategory | Birds | Bats |
| --- | --- | --- | --- | --- |
| Turbine Operation parameters | Reduce number of collisions | Blanket curtailment | None | Strong |
|  |  | Feathering | None | Strong |
|  |  | Algorithm-based curtailment | Limited | Limited |
|  |  | Activity-based curtailment | Limited | Limited |
| Altered lighting | Reduce attraction | Overall reduction of lighting | Limited | NA |
|  |  | Reduce use of white light | Strong | None |
|  |  | Alternative color choice | Mixed | Limited |
|  |  | Flashing lights | Strong | NA |
|  |  | Lighting intensity | None | NA |
| Turbine visibility/detectability enhancement | Increase visibility to reduce collisions | Adding flashing lights | Limited | NA |
|  |  | Painting turbine blades | Mixed | NA |
|  |  | Painting turbine towers | Limited | NA |
|  |  | UV light illumination | Limited | Mixed |
|  |  | Texturizing turbine blades | NA | None |
| Deterrents and other attraction reduction | Reduce attraction | Acoustic deterrents | Mixed | Limited |
|  |  | Visual deterrents | Mixed | NA |
|  |  | Physical deterrents | None | None |
|  |  | Chemical/olfactory deterrents | Mixed | NA |
|  |  | Combination deterrents | Limited | NA |
|  |  | Alter habitat/resources | Limited | Limited |
| Vessel operations | Reduce disturbance | Limit vessel speed | Limited | NA |
|  |  | Increase set-back distance | Limited | NA |

#### Turbine operation parameters

Altering turbine operation parameters has been a large focus of mitigation efforts in the terrestrial wind industry. Potential approaches (n=38) generally focused on reducing the amount of time turbines are spinning to reduce potential for collisions, with a large focus on curtailment, defined as preventing turbine blades from spinning during periods identified as high risk, often implemented via raising the wind speed threshold at which turbines begin to generate power (also called cut-in speed; Whitby et al. 2021). This approach broadly is known as blanket curtailment. “Feathering” also falls into this category, which is pitching turbine blades to catch as little wind as possible below cut-in speed, again reducing the time blades are spinning (Adams et al. 2021). Because reducing the amount of time turbine blades spin also leads to loss of energy production (Thurber et al. 2023), a range of approaches have been suggested for implementing targeted curtailment during periods of higher activity to maximize mortality reduction while minimizing energy loss, with variable levels of success. These include 1) algorithm-based curtailment during periods predicted to be of high risk defined based on environmental conditions (e.g., a combination of low wind speeds and higher temperatures; see Arnett and May 2016, Sinclair and DeGeorge 2016, American Wind Wildlife Institute 2019, Whitby et al. 2024), time of year (e.g., migration; Hill et al., 2014; Smallwood and Bell, 2020), and since this review was completed, incorporating both environmental conditions and bat acoustic activity patterns (Barré et al. 2023, Gottlieb et al. 2024, Peterson et al. 2025); and 2) activity-based curtailment that uses tools and technologies (e.g., radar, human observers, acoustic monitoring) to detect bats or birds within the vicinity of the turbine and trigger real-time turbine shutdown (Defingou et al. 2019).

While proposed for offshore wind, these approaches have only been tested in the terrestrial wind context. We found strong evidence in the literature that blanket curtailment was an effective mitigation strategy for bats in the terrestrial wind context. Multiple review papers suggested that across studies, a 50% reduction in bat mortality could be achieved by implementing cut-in speeds 1.5-3.0 m/s above manufacturer’s settings (Gartman et al. 2016, Adams et al. 2021, Whitby et al. 2021, 2024). For example, a meta-analysis of bat curtailment studies in the Midwest and Mid-Atlantic regions of the United States found that a 5.0 m/s cut-in speed could reduce total bat mortalities across facilities by an average of 62% (Whitby et al. 2021, 2024). There was also substantial evidence of effectiveness of feathering from multiple studies for bats at terrestrial wind facilities, either on its own or in combination with curtailment (Baerwald et al. 2009, Arnett and May 2016, American Wind Wildlife Institute 2019). There is also some evidence of effectiveness of algorithm-based curtailment for bats, particularly those based on both wind speed and temperature leading to a 62% reduction in bat mortality (Whitby et al. 2021) and activity-based real-time curtailment based on acoustic monitoring reduced bat mortalities by 83% (Hayes et al. 2019).

There is currently mixed evidence that curtailment reduces mortalities for birds. The primary focus to date in the terrestrial wind context has been raptors, with evidence for other bird groups generally lacking (Smallwood and Bell 2020). Studies with evidence of effectiveness for raptors utilized activity-based curtailment with human observers or automated detection systems at terrestrial wind farms to implement on-demand curtailment (Bennun et al. 2021, McClure et al. 2021, Ferrer et al. 2022). There has been debate in the scientific literature regarding the analytical approaches used in some cases to determine the reported degree of effectiveness (McClure et al. 2021, 2022, 2023, Huso and Dalthorp 2023).

Curtailment has been mentioned in the literature as a potential mitigation strategy in the offshore environment for both birds and bats, but this mitigation strategy has remained untested in this context. A review paper (Cook et al. 2011) suggested evidence of effectiveness for Marbled Murrelets (*Brachyramphus marmoratus*), but this was in relation to terrestrial wind farms (near nesting sites).

Variation in effectiveness may relate both to the ability to identify high risk periods during which to curtail, along with the degree to which a stationary turbine poses less risk than one that is spinning. Both factors may contribute to the apparent effectiveness of curtailment/feathering approaches for bats and the more mixed and taxon-specific results for birds.

#### Altered lighting to reduce attraction

There were 31 mitigation approaches in the reviewed literature related to reducing attraction via broadly limiting the use of lighting (n=18) as well as altering various lighting characteristics such as color (n=6), visibility (n=2), flashing (n=3), and intensity (n=2). Various approaches have been suggested to reduce light sources offshore, including reducing the use of flood lighting (Gauthreaux and Belser 2006, E-TWG Bird and Bat Specialist Committee 2020), reducing the number of turbines with lights (Mockrin and Gravenmier 2012), turning lights off when not in use, shielding lighting to only illuminate where needed (particularly down-shielding), and using on-demand lighting of various types (e.g., timers, heat sensors, motion sensors, ambient conditions, Aircraft Detection Lighting Systems; Gartman et al. 2016, E-TWG Bird and Bat Specialist Committee 2020). Many lighting minimization approaches have not been tested, with the exception of a few studies from islands that found lighting reduction and down-shielding decreased attraction of tubenose seabirds breeding in the vicinity (Reed et al. 1985, Miles et al. 2010). However, the use of flashing lights, different light intensity, and synchronicity of flashing have all been tested in various contexts. Flashing light was found to reduce avian attraction relative to continuously burning light in the terrestrial context (Evans et al. 2007, Gehring et al. 2009, Kerlinger et al. 2010, Cook et al. 2011, Gartman et al. 2016). One meta-analysis of avian collision mortality from 30 terrestrial wind farms across North America found no significant difference between mortality rates at turbines with red flashing aviation safety lighting and turbines without lighting at the same wind farms (Kerlinger et al. 2010). The more the flash duration was reduced, and the longer the time interval between flashes, the better for reducing wildlife interactions. However, there was no evidence of effectiveness of these measures offshore for birds, or any evidence of effectiveness for bats. In addition to lighting reduction, the color spectrum of artificial lights was also a focus of attraction minimization approaches. White light has been shown to attract birds more than other wavelengths of artificial light, but the color that leads to the least amount of attraction, is still debated. A study using lampposts in the Netherlands, for example, found that 60-81% of nocturnally migrating birds were disoriented and attracted by white lights, while fewer birds were influenced by red (54%), green (12-27%), or blue (4-5%) lights (Poot et al. 2008). However, a field test in the terrestrial United States that pointed lights into the night sky found that white, blue, and green lights all attracted birds, while red light did not (Evans et al. 2007). Based on field testing, several studies suggested the use of red light offshore, including near Procellarid breeding colonies (Syposz et al. 2021) and at offshore wind farms in the North Sea (Rebke et al. 2019). While the available evidence clearly indicated that limiting the use of white lights helped to mitigate attraction in birds, the best alternative lighting color is still unresolved, and may depend on weather, taxon, and other variables. More limited data was available on bat responses to light color. A study on the coast of the Baltic Sea found that bat acoustic activity increased around red LED light as opposed to white (Voigt et al. 2018) and increased with green light compared to no light (Voigt et al. 2017). However, high levels of bat activity around the white beams emitted by lighthouses has been noted for several decades (e.g., Cryan and Brown, 2007; Stantec Consulting Services Inc., 2016), and in a study of bat mortalities at a terrestrial wind farm in the United States, Bennett and Hale (2014) found that synchronized, flashing red aviation lights did not increase bat mortality rates compared to unlit turbines.

#### Turbine visibility and detectability enhancement

While in the instances described above, minimization methods were focused on reducing attraction to artificial light, there were some instances where the addition of lighting was proposed to increase visibility of turbines to reduce potential for bird or bat collisions (n=11). Many of these approaches have been field tested (n=6) or implemented (n=3), generally in terrestrial contexts. There is limited or mixed evidence for four bird-related approaches and one bat-related approach.

For birds, methods focused around adding lights and painting turbine blades to increase visibility. Day et al. (2005) determined that flashing lights served to encourage movement away from islands for eider species (*Somateria* sp.) in Alaska. Painting turbine blades was field tested and demonstrated to be effective for several bird species in the terrestrial wind context. An experimental laboratory study on American Kestrels (*Falco sparverius*) indicated that painting a single turbine blade black may reduce motion smear and could be effective for reducing bird collision risk (Hodos 2003). This strategy was further tested at a wind farm in Norway, in which May et al. (2020) found that painting one blade black significantly reduced the annual bird mortality rate by over 70% relative to neighboring unpainted turbines. Painting turbine foundations has also shown limited effectiveness for mitigating bird collisions at terrestrial wind facilities, with a 48.2% reduction in Willow Ptarmigan (*Lagopus lagopus*) mortality relative to unpainted turbine bases (Gartman et al. 2016, Stokke et al. 2020). As some birds have the ability to see in the ultraviolet (UV) spectrum, using UV-reflective paint to increase turbine visibility was also suggested but is untested in either onshore or offshore contexts. However, it is an established method for reducing bird strikes against windows (Marques et al. 2014, Defingou et al. 2019) and the use of UV light to illuminate electrical lines reduced avian collisions by 88% (Dwyer et al. 2019, Baasch et al. 2022). Previous studies suggested that UV light does not attract and disorient birds as much as longer wavelengths (e.g., as cited in Wiese et al. 2001), but additional data are needed on the utility of UV lights to illuminate obstacles and whether this is an effective approach to reduce potential avian collision risk offshore.

For bats, the primary focus of visibility and detectability enhancement is the use of UV light and painting or texturizing turbines. Gorresen et al., (2015) illuminated trees with dim flickering UV light (as an initial field test for use on turbines), which resulted in a reduction in Hawaiian hoary bat (*Lasiurus cinereus semotus*) activity levels despite an increase in insects (prey), though there was a high level of variation across sites. However, a western U.S. study of steady-burning dim UV light used to illuminate terrestrial wind turbines found no difference in bat mortality rates between illuminated and unilluminated turbines (Cryan et al. 2022). Other detectability enhancement measures have included texturizing turbine blades and towers to reduce attraction and mortality of bats; but field trials were inconclusive (Bennett and Hale 2018).

#### Deterrents and other attraction reduction

Acoustic, visual, physical and chemical deterrents and other (non-lighting related) approaches to minimize attraction were by far the most common category of mitigation approaches (n=38 approaches), with various methods suggested primarily in the terrestrial wind and agricultural contexts. Of these, 17 had not been tested or implemented, another 11 were field tested, and 10 were implemented. Of the 21 approaches tested, 14 showed limited or mixed evidence of effectiveness for birds and 4 showed limited evidence of effectiveness for bats. While some industries distinguish between deterrents and hazing devices, for the purposes of this review, we use ‘deterrents’ as an encompassing term to include both.

There are various types of deterrents aimed at scaring or otherwise repulsing individuals to counteract attraction and in turn reduce collision risk. In addition to acoustic (n=9), visual (n=7), physical (n=4), olfactory and chemical (n=2), combined method approaches (n=5), other types of approaches to reduce attraction relate to altering prey resources and habitat (n=11).

In the case of birds, acoustic deterrents include two types: bioacoustics, which primarily involve the playback of recorded alarm or distress calls by the species to be deterred; and other acoustic deterrents, which are generally loud noises used to scare birds. Both have been effectively used in agricultural and riparian settings (Enos et al. 2021), though effectiveness appears to vary by species, and habituation is common. Studies have shown that playing distress calls was an effective deterrent in agricultural contexts for both Black-crowned Night Herons (*Nycticorax nycticorax*) and Carrion Crow (*Corvus corone*) and for starlings in lab settings (Bomford and O’Brien 1990), though habituation was reported for starlings. In contrast, recorded alarm calls were ineffective at deterring Canada Geese (*Branta canadensis*) in recreational areas (Aguilera et al. 1991). Other studies have examined using loud noises (e.g., pyrotechnics, bangers, screamers, exploders) to deter birds from an area with varying effectiveness. For example, propane exploders reduced Red-winged Blackbird (*Agelaius phoeniceus*) damage to cornfields by 77%, with no evidence of habituation after 4 weeks (Conover 1984) and screamer shells reduced the number of geese in urban parks for up to 15 days (Aguilera et al. 1991). In contrast, response to pyrotechnics at a landfill site was species-specific, with a reduction in Herring Gull (*Larus argentatus*) numbers, but little response by crows and starlings (Curtis et al. 1993), and warning signals produced variable avoidance responses from eagles and other raptors at in terrestrial wind facility field tests (H.T. Harvey & Associates 2018, Felton et al. 2024). In general, bird habituation to loud noises may limit deterrent effectiveness over longer time periods. Presenting sounds at random intervals, using a range of different sounds or moving them frequently, and supporting sounds with additional auditory or visual methods can help to increase long-term effectiveness (Bomford and O’Brien 1990, Gilsdorf et al. 2002). In the offshore environment there is limited evidence regarding acoustic deterrents, though aquatic hazing devices were ineffective at deterring waterbirds species, such as Greater and Lesser Scaup (*Aythya affinis* and *A. marila*) and Surf Scoter (*Melanitta perspicillata*), from oil spills in the San Francisco Bay (Whisson and Takekawa 2000). Lastly, ultrasonic acoustic deterrents were implemented as a strategy to deter birds from perching at offshore platforms (Christensen-Dalsgaard et al. 2019), but there was minimal empirical proof of their effectiveness (Defingou et al. 2019).

Ultrasonic deterrents are the primary deterrent proposed for bats and have shown some evidence of effectiveness for reducing bat mortality at terrestrial wind facilities (O’Neil 2020, Hein et al. 2021) and riparian sites (Gilmour et al. 2020, 2021), though effectiveness may be location- and species-specific. For example, one study at a terrestrial wind farm found that broadband ultrasound decreased bat mortality between 2-64%, but only significantly for the hoary bat (*Lasiurus cinereus*) and silver-haired bat (*Lasionycteris noctivagans*; Arnett et al., 2013). Schirmacher (2020) did not detect a clear reduction in bat mortalities from the use of ultrasonic acoustic deterrents for any individual species, including hoary bats, silver-haired bats, big brown bats (*Eptesicus fuscus*), and eastern red bat (*Lasiurus borealis*). In fact, mortality of eastern red bat was estimated to be 1.3-4.2 times greater when deterrents were operational. Several studies tested the use of electromagnetic radiation deterrents on bats with some evidence of success (Nicholls and Racey 2007, 2009). Despite some promising evidence in the terrestrial environment, ultrasonic and electromagnetic deterrents have not yet been tested in the offshore context.

Visual deterrents are a common strategy used in agriculture and other terrestrial industries for birds (e.g., airports, landfills), and therefore many have been field tested and implemented in these settings. Visual deterrents have included lasers, effigies, flagging, flickering light, unmanned aerial vehicles (UAVs), and automated detection systems. Lasers have deterred certain bird species, including roosting Double-crested Cormorants (*Phalacrocorax auritus*) under low light conditions, but were not consistently effective across studies and did not disperse birds long distances (Glahn et al. 2000, Blackwell et al. 2002, Sherman and Barras 2004). In maritime industries, automated laser systems were successfully used on installations to deter gulls from large open areas such as helicopter landing platforms, and worked best in low light conditions (Christensen-Dalsgaard et al. 2019). Effigies, including scarecrows, hawk-kites, and eye-spot balloons, have also been used as a visual stimulus to deter birds in terrestrial contexts (Gilsdorf et al. 2002, Arnett and May 2016, Enos et al. 2021). For example, Conover (1984) reported plastic kites imprinted with a picture of a flying hawk reduced Red-winged Blackbird damage to cornfields by 83%, with no habituation observed during the 4-week study period and Stickley and King (1993) found that the use of a “Scary Man” pop-up inflatable effigy device proved effective in reducing Double-crested Cormorant numbers on catfish ponds for 10-19 days. There has been no testing of these devices in relation to OSW development. Reflective materials have also been tested as a visual deterrence method for birds in terrestrial contexts, with some success deterring Herring Gulls from nesting colonies, but not loafing sites (Belant and Ickes 1997). This approach was ineffective in reducing bird damage to agricultural fields, possibly due to habituation (Gilsdorf et al. 2002). Finally, UAVs, which include drones or remote-controlled helicopters, have been found to significantly reduce crop damage and bird abundances in several experiments on passerines in vineyards (Enos et al. 2021).

For bats, a 3-week study indicated that a UAV deterrent significantly reduced bat activity in the relevant airspace of up to 700m above the UAV’s flight altitude (Werber et al. 2023). There have not been any studies on the effectiveness of UAVs as deterrents in relation to terrestrial or offshore wind development. No other visual deterrents have been suggested or tested for bats (though see discussion regarding the use of dim UV light to reduce bat attraction, above).

Physical structural deterrents, such as wires, spikes, and netting, have been implemented as a method to mitigate avian perching or roosting behavior on terrestrial and offshore wind turbines and other infrastructure, but these approaches lacked evidence of effectiveness (Curry and Kerlinger 1998, Clarke 2004, Cook et al. 2011). Available research suggested that physical deterrents may work best when installed on horizontal structures (i.e., staircases, railings) and be more effective for certain species, including raptors and gulls (Christensen-Dalsgaard et al. 2019). No physical structural deterrents have been suggested for bats.

Several studies tested chemical deterrents (taste, olfactory, and tactile repellents) for birds at landfills, crop fields, and other terrestrial contexts with variable levels of effectiveness (Conover 1984, Curtis et al. 1993, Gilsdorf et al. 2002, Cook et al. 2011). In the offshore oil and gas industry, the use of ‘Bird Free Gel’, a sticky gel emitting UV light and with a deterring smell and taste, was implemented on an offshore oil installation in the North Sea and used successfully in other areas to deter breeding Black-legged Kittiwakes (*Rissa tridactyla*), gulls, and roosting pigeons (Christensen-Dalsgaard et al. 2019).

However, other research noted that permanence of chemical repellents may vary depending on conditions such as terrain and weather conditions (Marques et al. 2014, May et al. 2015, Arnett and May 2016). In general, there was little evidence of effectiveness for any of these deterrents in the offshore wind context, and the level of effectiveness varied based on species, environmental conditions, and other factors. Again, this minimization category was not proposed or applied in relation to bats.

Several studies assessed integration of multiple deterrent techniques for birds. Effective examples included using multiple acoustic deterrents to reduce abundance of Laughing Gulls (*Leucophaeus atricilla*) at an airport (Montoney and Boggs 1993) and Canada Geese on lawns (Mott and Timbrook 1988), and combining acoustic and chemical deterrents to reduce waterfowl landings at contaminated power plant pools (Stevens et al. 2000). However, a pilot study at a terrestrial wind facility in Canada determined that using predator owl deterrent models and bioacoustic alarms did not significantly deter birds from wind turbines (Dorey et al. 2019). As such, the effectiveness of integrated deterrent approaches for birds at terrestrial wind facilities remains variable, and in our review of the literature, no similar approaches were tested at OSW facilities.

Lastly, efforts to reduce bird attraction to certain structures or areas have used habitat enhancement in other locations. In the terrestrial wind context, it has been suggested that supplementary or diversionary feeding could be effective in reorienting raptors away from wind farms (Simmons et al. 2020). However, while a study of Hen Harriers (*Circus cyaneus*) showed an increase of 32-42% in flight activity in a created habitat enhancement area next to a terrestrial wind facility, there was no difference in activity within the wind facility (Gartman et al. 2016). In contrast, the felling of a plantation forest increased Golden Eagle use of the enhanced area, with a shift in their range away from the terrestrial wind farm (Gartman et al. 2016). Reducing the attractiveness of the wind farm itself has also been tested in the terrestrial wind context with some effectiveness, including reducing attraction for Lesser Kestrels (*Falco naumanni)* by tilling soil at the base of turbines (Garcia-Rosa and Tande 2023) or removing rock piles attractive to rodent prey (Thelander et al. 2003). Decoy towers without functioning rotor blades, positioned at the end of turbine rows, were also tested as a potential method to deter raptor species from perching at an terrestrial wind facility, but effectiveness was not determined (Curry and Kerlinger 1998) and it is unclear the degree to which this might increase attraction to the general wind farm area (Smallwood and Karas 2009, Cook et al. 2011).

#### Vessel operations

While the above minimization methods focus on reducing collisions and/or attraction, there are additional operational considerations specific to marine industries focused on vessels (n=6 approaches), primarily relevant to marine birds with the goal of reducing disturbance. Approaches included those aimed at reducing bird disturbance by minimizing vessel speed and density, implementing setbacks from high-use areas (i.e., breeding or foraging grounds), and using caution in low visibility conditions. Two approaches, limiting vessel speed and volume and adopting set-back distances, were field tested as mitigation strategies in maritime industries with some evidence of effectiveness. For example, Bellefleur et al. (2009) determined that Marbled Murrelets required a minimum buffer distance of 29 m between a boat and birds on the water to reduce disturbance such that 75% of the studied population would be minimally affected. Additionally, Ronconi and St. Clair (2002) found that using a vessel setback distance of 600 m from shore with a speed limit of 25 km/h reduced Black Guillemot (*Cepphus grylle*) flushing behavior to ∼10%. No studies examined the effectiveness of implementing vessel operation parameters for OSW development specifically..

### Compensation

Compensation for environmental effects that cannot be avoided or minimized may occur on or off site and may include both in-kind or out-of-kind compensation (e.g., providing a resource of similar or different structural or functional type to the affected resource). While compensatory mitigation is in many cases implemented as part of permitting, it can also be pursued voluntarily. Compensation-related mitigation approaches (n=24) included those focused on the methods and funding of such approaches (n=10), as well as different types of compensation implementation projects (n=14).

Compensation methods and funding approaches included mitigation banking and in-lieu fee programs (Arnett and May 2016). A mitigation bank is a site or suite of sites that provides ecological functions and services expressed as credits that are conserved and managed in perpetuity for particular species and are used expressly to offset effects occurring elsewhere to the same species. In-lieu fee programs, which are similar to mitigation banks, provide ecological functions and services expressed as credits that are conserved and managed for particular species or habitats and used to offset effects occurring elsewhere to the same species or habitats (USFWS, 2023). Mitigation banks have been implemented in some cases in the terrestrial wind industry as well as other projects that require habitat restoration (Arnett and May 2016). Other methods suggested include development of in-lieu fees and funding pools, permittee-required compensation strategies, financial support of rehabilitation programs, and approaches to calculate offsets (Orloff and Flannery 1992, Cole 2011, Levrel et al. 2012, Ronconi et al. 2015, Millon et al. 2021), though these had not been implemented in the source literature.

Compensation implementation approaches for birds primarily focused on increasing quality habitat in other areas via habitat creation and restoration (Marques et al. 2014, Arnett and May 2016, McGregor et al. 2022, Liang et al. 2023), restriction of access (either for fishing or recreation; Lüdeke 2017), and reducing other sources of mortality via invasive species management, reduction of bycatch, or other methods. One strategy that has been extensively tested focused on the removal of mammalian predators to increase colonial seabird demographics. Removal of invasive predators from breeding colonies has generally been found to increase productivity (e.g., Pascal et al. 2008, Nogales et al. 2013) though with some variation in success by species. For example, Luxmoore et al. (2019) found that Guillemot/Murre (*Uria* spp.) populations did not respond to the removal of rats, while Booker et al. (2019) attributed increases in closely-related Razorbill (*Alca torda*) and Atlantic Puffin (*Fratercula arctica*) breeding numbers to the removal of rats (Booker et al. 2019). Thus, the effectiveness of these types of strategies is likely context-specific.

In the case of bats, compensation implementation has primarily been habitat-based in the terrestrial wind context to compensate for collision mortalities, and there was some evidence of effectiveness, including increased bat species presence with the creation of natural features (e.g., hedgerows, fallows) in agricultural areas with terrestrial wind developments (Millon et al. 2015) and increased bat abundance with the creation of water resources and roost boxes (Brack et al. 2022).

## DISCUSSION

In our literature review, we found few mitigation approaches available for birds and bats that have been adequately tested, particularly with regards to the OSW industry. We lack evidence of effectiveness for a vast majority of the mitigation approaches identified (birds: 81%; bats 89%).

Commonly implemented approaches often had not been demonstrated to be effective (e.g., raising the lower end of the rotor swept zone to reduce marine bird collision risk) or had been shown to be effective in other scenarios such as with terrestrial wind energy development, but not tested offshore (e.g., curtailment of turbine operations based on environmental conditions to reduce bat collision risk). For birds, approaches related to lighting (reducing artificial light, avoiding white and steady-burning lights, etc.) were the most commonly tested and effective methods for reducing maladaptive attraction and collisions. For bats, alterations of turbine operations (e.g., curtailment and feathering of turbine blades) were most commonly shown to be effective. Most minimization approaches have primarily been tested in terrestrial contexts, however, leaving important questions regarding the transferability of these results to the offshore environment. This lack of evidence in the offshore environment points to the challenges associated with this type of assessment, including a lack of standard definitions and metrics, a lack of fundamental understanding of the mechanisms that may influence animal responses to offshore wind energy development or to mitigation approaches, and the need to explore effectiveness of different types of mitigation at different spatial (and sometimes temporal) scales and across relevant taxa with different morphological and sensory characteristics.

### Challenges with Assessing Effectiveness

There is no widely accepted definition of “effectiveness” for OSW mitigation approaches nor standardized metrics with which use in these assessments, which led to challenges for the purposes of this review. One recent assessment (Crown Estate Scotland 2024) defined mitigation success as situations in which “no deleterious effects were recorded,” but the purpose of mitigation is to avoid, minimize, or compensate for impacts that might otherwise occur, not to avoid creating further impacts. Another approach, the Conservation Evidence Project, which is more broadly focused on conservation methods to maintain and restore biodiversity, uses expert elicitation to assess the effectiveness of specific conservation interventions and the uncertainty associated with the available evidence (Sutherland et al. 2021). While a standardized framework is used for integrating expert input, assessments of effectiveness are made relative to the specific conservation objectives and outcomes that each intervention is intended to achieve (Sutherland et al. 2021). In the offshore wind context, an expert elicitation-style approach to evaluating evidence and associated uncertainty could be a valuable method for determining which mitigation approaches have sufficient evidence and certainty to be implemented at OSW facilities.

Standard definitions and clearly articulated goals would be helpful to inform research efforts to quantitatively assess effectiveness of mitigation approaches for birds and bats in the offshore wind context, but need to consider the category of the mitigation hierarchy and the effect that is being mitigated (e.g., the definition of effectiveness may vary for a compensation measure vs. a minimization measure).

Challenges in assessing mitigation effectiveness in the OSW context arise from a combination of limitations in measurement and metrics, uncertainty regarding underlying biological mechanisms, and logistical constraints associated with conducting studies in offshore environments. Monitoring programs to assess mitigation effectiveness can be difficult to design and implement, particularly in the offshore context. Studies must be designed with sufficient statistical power, including acquisition of enough data to reliably detect a change in the selected effect metric if it occurs (Martins et al. 2023, Regional Synthesis Workgroup of the Environmental Technical Working Group 2023). Approaches for assessing the effectiveness of mitigation approaches will also vary based on whether they fall within the avoidance, minimization, or compensation categories of the mitigation hierarchy.

It is generally difficult to test the effectiveness of avoidance measures, because there are typically no within-site “control” scenarios against which impacts can be compared. As a result, effectiveness of avoidance approaches must primarily be inferred from natural experiments, for example by comparing outcomes across projects sited in locations that vary in wildlife value. Similarly, assessing the effectiveness of approaches implemented during the project design phase (e.g., changes to turbine height, rotor diameter, air gap, turbine spacing or layout, etc.) requires comparisons across multiple wind facilities with differing characteristics to evaluate their influence on effects. Evaluating effectiveness in these cases is only possible when data collection methods are harmonized across projects and data are publicly available, enabling robust multi-facility analyses.

In comparison, minimization approaches, such as use of deterrents or changes to turbine operations, are generally more straightforward to assess using standard control-impact study design (Smokorowski and Randall 2017), for example by implementing the mitigation approaches at a subset of turbines within a single wind facility. Even for these approaches, however, standardized metrics and public data availability remain essential to enable comparisons across projects and taxa as meta-analyses are often required for statistical power, and technological advancements in remote collisions detection will be important for improving the ability to evaluate effectiveness specifically for approaches aimed at minimizing collisions (Skov et al. 2025).

In contrast, compensatory mitigation approaches often have broader goals and are frequently implemented away from the impacted location, which complicates assessments of effectiveness. Compensation approaches may include a wide range of interventions operating at species, community or ecosystem levels to offset ecological loss (Thébaud et al. 2015). In the context of birds and bats and OSW development, the primary objective of compensation is to offset mortality by increasing population size or fitness. Assessing effectiveness therefore requires an understanding of population-level constraints, including other anthropogenic stressors, density dependence, and metapopulation dynamics, as well as the ability to accurately measure demographic rates and population size.

Finally, much of the evidence base reviewed in this study comes from outside of the OSW industry, raising the question about the transferability of effectiveness across contexts. While extrapolation across taxa, systems and spatiotemporal settings is common in ecology (Houlahan et al. 2017), exposure pathways and risk profiles differ substantially between offshore and terrestrial wind development. For bats, offshore exposure is thought to be largely limited to migratory tree bats (Solick and Newman 2021), representing a subset of species affected by terrestrial wind developments (Thompson et al. 2017), and uncertainty remains regarding both collision risk and the mechanisms driving offshore exposure (Hein et al. 2021, Guest et al. 2022). For birds, species exposed offshore differ markedly from those targeted by terrestrial wind farm mitigation approaches, further limiting transferability. Our review indicates that mitigation effectiveness is likely to be species- and location-specific, and approaches shown to be effective in other contexts should be tested for OSW applications and focal species where possible. Given these constraints, developing and testing a range of mitigation options, supported by improved data quality and availability, will be critical for evaluating transferability and effectiveness in offshore settings (Yates et al. 2018).

### Implementation and Testing of Mitigation Approaches

Implementation of unproven mitigations creates the assumption that effects to birds and bats are being adequately addressed when that may not be the case. Given the general lack of strong evidence for the effectiveness of mitigation approaches in the OSW context, we echo other recommendations in relation to wind energy development (e.g., May 2017) and encourage 1) avoidance approaches as a key first step in the mitigation hierarchy (Council on Environmental Quality 2020), considered by many to be the most certain way of managing negative impacts to wildlife (Phalan et al. 2018, Bennun et al. 2021), 2) prioritizing implementation of well-tested minimization measures with reasonable evidence of effectiveness, should the level of risk warrant the implementation, 3) testing of other promising minimization approaches for which data are currently lacking (this must include the development of standardized assessment metrics to determine mitigation effectiveness), and 4) exploring compensation approaches for effects that cannot currently be reliably avoided or minimized.

There are a few key mitigation approaches that could likely be implemented across offshore wind facilities without the need for further study and testing, due to their place within the mitigation hierarchy and/or a strong base of evidence from other industries. Other approaches may be promising, with limited evidence or in some cases conflicting evidence, and thus require additional research before widespread offshore implementation. This does not imply that these mitigation approaches should be prohibited, but rather that they should not be credited as effective impact reduction without adequate evidence. Where such approaches are applied, they should be framed explicitly as experimental, with clear hypotheses, performance metrics, and robust follow-up monitoring to evaluate whether they reduce mortality or population-level risk from OSW development. In addition to the existing body of evidence, research on the effectiveness of mitigation approaches should consider 1) the degree to which a mitigation approach has already been implemented at offshore wind farms (even without evidence of effectiveness), as those already included in project permits are also more likely to be proposed at new projects, thus making them a higher priority for further examination, 2) our current understanding of risk (e.g., prioritizing measures with the greatest potential to mitigate effects), 3) the scale of research needed to assess effectiveness (studies that require comparisons across projects with different project designs will necessarily require the construction of multiple projects with differing characteristics and longer-term data collection to facilitate comparisons compared to those that can be tested at individual projects), and 4) our current ability to assess effectiveness (in particular, technological advancements are needed to better measure the effectiveness of minimization measures focused on reducing collisions in the offshore context).

#### Avoidance

There is a strong scientific consensus that siting and other avoidance measures remain a key first step to reduce OSW development impacts, particularly for birds (Langston and Pullan 2003, Phalan et al. 2018, Fox and Petersen 2019, Danovaro et al. 2024). As described above, there are many challenges hindering our ability to assess effectiveness of avoidance measures, and thus we generally recommend implementation even with limited evidence for effectiveness as a first step in the mitigation hierarchy, but with careful consideration and monitoring.

We also suggest prioritizing understanding the influence of adjustments to turbine size and air gap because this is an avoidance approach that has already been implemented at OSW farms and included in project permits in jurisdictions worldwide (e.g., Crown Estate Scotland 2024), making it more likely to be proposed at new projects. The specifications of turbines (including size and air gap) may influence both how animals perceive the wind farm as a whole and how they utilize space in and around the wind farm. While there is some evidence of the importance of turbine size and air gap for reducing collisions in the terrestrial wind context (Garvin et al. 2024), a great deal of additional research is needed to understand the degree to which these factors affect birds and bats in the offshore context and the factors influencing variability in results. As with avoidance, this type of multi-facility study requires development activities to proceed at a range of locations (such that projects with different characteristics can be compared), along with consistent data collection methods across projects and public availability of data from site-specific studies. We suggest that where such approaches are implemented, they be accompanied by a rigorous regional-scale monitoring plan to assess their effectiveness.

#### Minimization

##### Reductions in artificial light

The use of alternative lighting options to reduce effects to birds is perhaps the best-understood and most thoroughly tested mitigation category we reviewed. Artificial lighting can have serious impacts on wildlife populations, species, and ecosystems (Jägerbrand and Bouroussis 2021, Walsh et al. 2024), therefore we propose all OSW facilities enact mitigation to reduce artificial light pollution and avoid attracting and disorienting birds, particularly nocturnal migrants. This can be achieved by minimizing the use of steady-burning and white lights whenever possible, down-shielding of lights, reduction of flood lighting during construction and maintenance, and other approaches detailed in guidance documents for OSW development (E-TWG Bird and Bat Specialist Committee 2020, BOEM 2021) as well as other industries (e.g., Environment and Climate Change Canada 2016). While lights are required on turbines and other offshore infrastructure for aviation and marine navigation safety purposes, and regulations vary by country, it is recommended that the use of aircraft detection lighting systems and similar mechanisms to reduce lighting at night be implemented wherever possible, including adjustments of current regulations, where needed, to allow for such measures. Based on evidence from this review, use of non-white, flashing, less intense, shielded lights - with those lights operating as infrequently as possible - is the best available approach to reduce avian attraction, disorientation, and mortality around offshore wind structures. Nevertheless, additional research would be beneficial to better understand which wavelengths of (non-white) light can best reduce risk for different weather conditions and taxa, particularly in the offshore environment, since available findings are contradictory. A better understanding of bat response to light would also be useful, as uncertainty remains regarding the mechanisms of attraction for this taxon (Guest et al. 2022). Our relatively poor understanding of animals’ vision likely hinders our ability to make informed decisions about mitigation in this area.

##### Increasing visibility of turbine blades

The use of paint for increasing visibility of turbines and other structures shows promising evidence but has to date been tested infrequently and with few species (Hodos 2003, May et al. 2020). There is the most consistent evidence of effectiveness for raptors; with recent laboratory evidence for European bats (Jonasson et al. 2025), while use of this approach with marine birds requires additional study. Additional study of bird and bat responses to UV light could also help inform its use as a possible mitigation approach. Differences in vision among bird (and bat) taxa suggest that vision-related approaches to enhance turbine visibility must be carefully considered and tested.

##### Deterrents

In many cases the studies of effectiveness of deterrents focused on one or a few species over a short period of time and often showed variable results. While deterrents have been used in the OSW industry in Europe to reduce perching for taxa such as gulls and cormorants (Gartman et al. 2016), these facilities were located closer to shore than those being built to date in the U.S. Atlantic. Consequently, the level of attraction (for roosting, perching, and nesting) is site-specific and may vary with distance from shore and therefore implementation should be tailored to the site. As discussed above, the use of OSW turbines by bats remains largely unknown in the region as well, though studies from Europe have found that bats may roost on OSW turbines (Ahlen et al. 2009, Lagerveld et al. 2014) with uncertainty in the frequency of occurrence and connection to collision risk. Thus, for this type of mitigation, we recommend not only additional research on effectiveness, but also to consider implementation in situations where there is evidence that attraction is occurring at large enough scales to warrant investment in deterrents. While in general it is thought that reducing perching and roosting opportunities can help reduce attractiveness of wind farms to birds, which in turn may help reduce collision risk (Gartman et al. 2016), we lack direct evidence linking perching deterrents to collision risk reduction. Key studies to fill knowledge gaps include those focused on species that occur offshore and long-term studies to understand the potential for habituation to deterrents in that environment.

##### Turbine Operational Parameters

One of the largest data gaps identified in this review (as well as others; Hein et al. 2021) is the degree to which bat mortality at offshore wind farms needs to be mitigated. Feathering and curtailment have been shown to be effective at reducing collision for bats in the terrestrial context. While there is evidence that migratory tree roosting bats occur offshore, uncertainty remains in both the abundance and distribution of bats in the offshore environment as well as the degree to which collision risk and associated mortality mirrors patterns onshore (Solick and Newman 2021). However, there may be instances that warrant implementation without further testing where the level of risk based on current knowledge and potential for population-level effects if unmitigated are high. Additional research is also needed for birds before this type of approach is implemented in the offshore context targeted at birds. The available evidence suggests that curtailment based on wind speed is ineffective for reducing avian mortality at terrestrial wind farms (e.g., Smallwood and Bell 2020). There is some evidence that activity-based curtailment may be effective for particular species, primarily raptors (e.g., when they have been detected entering a wind farm), but the drivers of raptor risk and mortality may differ from other taxa such as marine birds. While there have been recent advances in radar and video systems to detect birds and bats at OSW farms, perhaps the greatest challenge to testing these types of strategies relates to the difficulty in collecting adequate data on collision rates (e.g., sample size across rotor sweep zone and wind facility). The further development and testing of collision detection technologies represents a key step towards better understanding mortality risk for bats and birds at OSW facilities to in turn inform the utility of curtailment in this context.

##### Minimizing vessel disturbance

Despite the small number of sources and mitigation approaches focused on this topic, behavioral disturbance is a common effect from OSW development for marine birds (Allison et al. 2019, Fox and Petersen 2019, Williams et al. 2024). In addition to responses to the turbines themselves, birds may be disturbed due to vessel and other OSW-related activity, which has the potential to have similar energetic consequences. Altering vessel speed and approach distance to wildlife are key mitigation approaches for marine mammals and sea turtles for the OSW and other maritime industries (Hazel et al. 2007, Conn and Silber 2013). These strategies have the potential to benefit marine birds as well. While the impact of this type of disturbance is not well studied, it may be an important mitigation strategy to evaluate given its potential effectiveness in the maritime environment. Thus, additional research to understand variation in response among bird species, implementation of best practices, and the potential consequences of vessel disturbance would go a long way to understand the degree to which this type of effect could be mitigated effectively for marine birds.

#### Compensation

Without on-site mitigation approaches that have been proven to adequately ameliorate effects of development, off-site and potentially out-of-kind measures (e.g., via compensation) must be considered in order to address remaining unmitigated effects (Croll et al. 2022). In the case of birds, most compensatory mitigation has focused on marine birds, and included approaches related to fisheries (e.g., fisheries closures for species that are key prey for seabirds, efforts to reduce fisheries bycatch), creation or restoration of nesting habitat and marine reserves, and the eradication and control of predators, including invasive mammals like rats (Jodice et al. 2019, McGregor et al. 2022). For bats, compensation often involves habitat creation and restoration.

Understanding the effectiveness of these types of approaches is challenging, as it generally requires long-term monitoring of populations to examine changes in key demographic parameters along with a quantitative understanding of the influences of other anthropogenic threats on these populations. While in some cases there may be no well-established offset approach that is known to be effective for a given species, there is clear evidence that colony-based and island-based mitigation approaches to control invasive predators (in the case of seabirds, for example), do increase breeding success (e.g., Pascal et al. 2008, Nogales et al. 2013). A recent review focused on OSW development in the United States noted that compensatory mitigation may be the best approach to adequately prevent cumulative impacts that lead to population declines across both regional and international scales, and that implementation will in turn require a strategically coordinated approach (Croll et al. 2022). As with avoidance and minimization, compensatory mitigation must be carefully designed and monitored to assess effectiveness in mitigating OSW effects.

## CONCLUSIONS

Few offshore wind energy mitigation approaches for birds and bats have been shown to be effective, due in part to challenges with defining effectiveness of mitigation and a lack of standardized methods, as well as challenges associated with conducting tests of such measures in offshore environments. However, there are several avenues of future research that could be prioritized for immediate investigation. First, we recommend that research focus on mitigation approaches that have already been implemented at offshore wind farms and included in project permits in jurisdictions worldwide. This is important since approaches that have already been implemented at existing projects are more likely to also be proposed at new projects. This includes project design measures such as increasing the air gap to reduce collisions of marine birds. Second, we recommend focusing research on mitigation approaches with the greatest potential impact in reducing effects, particularly those focused on reducing direct mortality. For example, curtailment and feathering for bats falls into this category given the proven effectiveness and prevalence of this impact at terrestrial wind farms Adams et al. 2021, Whitby et al. 2024), with research focused on understanding the level and factors affecting exposure.

Curtailment for birds is a commonly discussed measure in the offshore wind context with little evidence of effectiveness (apart from activity-based curtailment) that could also fall into this category. Thirdly, we recommend an immediate focus on mitigation approaches that can be tested at the individual project level, such as visibility enhancement measures. Studies that require comparisons across projects with different project designs will necessarily require the construction of multiple projects with differing characteristics and longer-term data collection to facilitate comparisons; many of these measures may have the greatest potential to reduce effects but will be difficult to adequately test in the immediate term. Conducting well-designed studies of effects to birds and bats from OSW development, as well as ensuring the public availability of resulting effects data, is an essential step to allow for these longer-term studies of the influence of avoidance measures on wildlife mitigation. And fourthly, additional technology development may be needed in some cases to assess mitigation effectiveness. In particular, technological advancements are needed to better measure the effectiveness of minimization measures focused on reducing collisions in the offshore context.

## ACKNOWLEDGEMENTS

This project was funded by the New York State Energy Research and Development Authority (NYSERDA) through support for the Mitigation Practices Database, and by Environment and Climate Change Canada’s Wildlife and Landscape Science Directorate (Grants and Contributions). We thank members of the Offshore Wind Environmental Technical Working Group (E-TWG) for oversight and input, and NYSERDA project manager K. McClellan Press. We also thank I. Stenhouse and L. LaMartina for project support; J. Wisbey and A. Millard for citation management; Sylvain Christin for French translation; and Rush Dhillon (Rush Studio) for design of figures and the graphical abstract.

This review was completed by authors based in Canada and the United States, and we acknowledge with respect the Indigenous Peoples on whose lands and waters we live and work. This includes the Wabanaki Peoples (Mi’kmaq, Wolastoqiyik/Maliseet, Peskotomuhkati/Passamaquoddy, Penobscot), the Beothuk, and the Anishinàbe Algonquin People. We also acknowledge that the research synthesized draws on studies conducted across many regions, and therefore across the lands and waters of Indigenous Peoples around the world.

## DATA AVAILABILITY

This study is a review of previously published literature. No new datasets were generated or analyzed. References for all data are included in Appendices A-B. The data that support the findings of this literature synthesis study are openly available in Mitigation Practices Database Tool at (https://briloon.shinyapps.io/MPDTool/).

**APPENDIX A. Mitigation Review References (CSV)**

**APPENDIX B. Mitigation Review References (BibTeX)**

## REFERENCES

Adams, E. M., J. Gulka, and K. A. Williams. 2021. A review of the effectiveness of operational curtailment for reducing bat fatalities at terrestrial wind farms in North America. I. Torre, editor. PLOS ONE 16:e0256382.

Aguilera, E., R. L. Knight, and J. L. Cummings. 1991. An Evaluation of Two Hazing Methods for Urban Canada Geese. Wildlife Society Bulletin 19:32–35.

Ahlen, I., H. J. Baagøe, and L. Bach. 2009. Behavior of Scandinavian Bats during Migration and Foraging at Sea. Journal of Mammalogy 90:1318–1323.

Allison, T. D., J. E. Diffendorfer, E. F. Baerwald, J. A. Beston, D. Drake, A. M. Hale, C. D. Hein, M. M. Huso, S. R. Loss, J. E. Lovich, M. D. Strickland, K. A. Williams, and V. L. Winder. 2019. Impacts to wildlife of wind energy siting and operation in the United States. Issues in Ecology.

American Wind Wildlife Institute. 2019. AWWI Technical Report: A Summary of Bird Fatality Data in a Nationwide Database. Washington, DC. Available at www.awwi.org.

Arnett, E. B., and R. F. May. 2016. Mitigating wind energy impacts on wildlife: approaches for multiple taxa. Human-Wildlife Interactions 10:28–41.

Arnett, E., C. Hein, M. Schirmacher, M. Huso, and J. Szewczak. 2013. Correction: evaluating the effectiveness of an ultrasonic acoustic deterrent for reducing bat fatalities at wind turbines. PLOS ONE 8: 10.1371/annotation/a81f59cb-0f82-4c84-a743-895acb4.

Baasch, D. M., A. M. Hegg, J. F. Dwyer, A. J. Caven, W. E. Taddicken, C. A. Worley, A. H. Medaries, C. G. Wagner, P. G. Dunbar, and N. D. Mittman. 2022. Mitigating avian collisions with power lines through illumination with ultraviolet light. Avian Conservation and Ecology 17.

Baerwald, E. F., and R. M. R. Barclay. 2011. Patterns of activity and fatality of migratory bats at a wind energy facility in Alberta, Canada. Journal of Wildlife Management 75:1103–1114.

Baerwald, E. F., J. Edworthy, M. Holder, and R. M. R. Barclay. 2009. A Large-Scale Mitigation Experiment to Reduce Bat Fatalities at Wind Energy Facilities. Journal of Wildlife Management 73:1077–1081.

Barré, K., J. S. P. Froidevaux, A. Sotillo, C. Roemer, and C. Kerbiriou. 2023. Drivers of bat activity at wind turbines advocate for mitigating bat exposure using multicriteria algorithm-based curtailment. Science of The Total Environment 866:161404.

Belant, J. L., and S. K. Ickes. 1997. Mylar Flags as Gull Deterrents. Pages 73–80 in. Proceedings of the Thirteenth Great Plains Wildlife Damage Control Workshop.

Bellefleur, D., P. Lee, and R. A. Ronconi. 2009. The impact of recreational boat traffic on Marbled Murrelets (Brachyramphus marmoratus). Journal of Environmental Management 90:531–538.

Bennett, V., and A. Hale. 2018. Resource Availability May Not Be a Useful Predictor of Migratory Bat Fatalities or Activity at Wind Turbines. Diversity 10:44.

Bennett, V. J., and A. M. Hale. 2014. Red aviation lights on wind turbines do not increase bat-turbine collisions. Animal Conservation 17:354–358.

Bennun, L., J. Van Bochove, C. Ng, C. Fletcher, D. Wilson, N. Phair, and G. Carbone. 2021. Mitigating biodiversity impacts associated with solar and wind energy development: guidelines for project developers. IUCN, The Biodiversity Consultancy, Gland, Switzerland; Cambridge, UK.

Blackwell, B. F., G. E. Bernhardt, and R. A. Dolbeer. 2002. Lasers at Nonlethal Avian Repellents. The Journal of Wildlife Management 66:250–258.

BOEM. 2021. Guidelines for Lighting and Marking of Structures Supporting Renewable Energy Development. United States Department of the Interior Bureau of Ocean Energy Management Office of Renewable Energy Programs, Stirling, VA.

Bomford, M., and P. H. O’Brien. 1990. Sonic Deterrents in Animal Damage Control: A Review of Device Tests and Effectiveness. Wildlife Society Bulletin 18:411–422.

Brack, V. J., D. W. Sparks, and S. Kennedy. 2022. Case Study: Upland Ponds Provide On-Site Mitigation for Bat Habitat along American Electric Power’s 765-kV Powerline ROW in the Appalachian Mountains, USA. L. Hufnagel, editor. New Insights Into Protected Area Management and Conservation Biology. IntechOpen.

Burt, C. S., J. F. Kelly, G. E. Trankina, C. L. Silva, A. Khalighifar, H. C. Jenkins-Smith, A. S. Fox, K. M. Fristrup, and K. G. Horton. 2023. The effects of light pollution on migratory animal behavior. Trends in Ecology & Evolution 38:355–368.

Christensen-Dalsgaard, S., N. Dehnhard, B. Moe, G. H. R. Systad, and A. Follestad. 2019. Unmanned installations and birds. A desktop study on how to minimize area of conflict. Norwegian Institute for Nature Research., Trondheim, Norway.

Cohen, E. B., J. J. Buler, K. G. Horton, S. R. Loss, S. A. Cabrera-Cruz, J. A. Smolinsky, and P. P. Marra. 2022. Using weather radar to help minimize wind energy impacts on nocturnally migrating birds. Conservation Letters 15:e12887.

Cole, S. 2011.How much is enough? Adequate Levels of Environmental Compensation for Wind Power Impacts Using Equivalency Analysis: An Illustrative & Hypothetical Case Study of Sea Eagle Impacts. Swedish Agricultural Institution.

Conn, P. B., and G. K. Silber. 2013. Vessel speed restrictions reduce risk of collision-related mortality for North Atlantic right whales. Ecosphere 4:1–16.

Conover, M. R. 1984. Comparative Effectiveness of Avitrol, Exploders, and Hawk-Kites in Reducing Blackbird Damage to Corn. The Journal of Wildlife Management 48:109–116.

Cook, A. S. C. P., E. M. Humphreys, F. Bennet, E. A. Masden, and N. H. K. Burton. 2018. Quantifying avian avoidance of offshore wind turbines: Current evidence and key knowledge gaps. Marine Environmental Research 140:278–288.

Cook, A. S. C. P., V. H. Ross-Smith, S. Roos, N. H. K. Burton, N. Beale, C. Coleman, H. Daniel, S. Fitzpatrick, E. Rankin, K. Norman, and G. Martin. 2011. Identifying a range of options to prevent or reduce avian collision with offshore wind farms using a UK-based case study. British Trust for Ornithology, Thetford, UK.

Council on Environmental Quality. 2020. Protection of the environment (under the National Environment Policy Act). Council on Environmental Quality, Washington, D.C.

Crawford, R. L., and R. T. Engstrom. 2001. Characteristics of Avian Mortality at a North Florida Television tower: a 29-Year Study. Journal of Field Ornithology 72:380–388.

Croll, D. A., A. A. Ellis, J. Adams, A. S. C. P. Cook, S. Garthe, M. W. Goodale, C. S. Hall, E. Hazen, B. S. Keitt, E. C. Kelsey, J. B. Leirness, D. E. Lyons, M. W. McKown, A. Potiek, K. R. Searle, F. H. Soudijn, R. C. Rockwood, B. R. Tershy, M. Tinker, E. A. VanderWerf, K. A. Williams, L. Young, and K. Zilliacus. 2022. Framework for assessing and mitigating the impacts of offshore wind energy development on marine birds. Biological Conservation 276:109795.

Crown Estate Scotland. 2024. Collaboration for Environmental Mitigation & Nature Inclusive Design (CEMNID) Mitigation Measures Efficacy Review. Xodus Group.

Cryan, P. M., and A. C. Brown. 2007. Migration of bats past a remote island offers clues toward the problem of bat fatalities at wind turbines. Biological Conservation 139:1–11.

Cryan, P. M., P. M. Gorresen, B. R. Straw, S. (Simon) Thao, and E. DeGeorge. 2022. Influencing Activity of Bats by Dimly Lighting Wind Turbine Surfaces with Ultraviolet Light. Animals 12:9.

Curtis, P. D., C. R. Smith, and W. Evans. 1993. Techniques for reducing bird use at Nanticoke landfill near E.A. Link Airport, Broome County, New York. Page 11 in. Sixth Eastern Wildlife Damage Control Conference.

Danovaro, R., S. Bianchelli, P. Brambilla, G. Brussa, C. Corinaldesi, A. Del Borghi, A. Dell’Anno, S. Fraschetti, S. Greco, M. Grosso, E. Nepote, L. Rigamonti, and F. Boero. 2024. Making eco-sustainable floating offshore wind farms: Siting, mitigations, and compensations. Renewable and Sustainable Energy Reviews 197:114386.

Defingou, M., F. Bils, B. Horchler, T. Liesenjohann, and G. Nehls. 2019.PHAROS4MPAs: A Review of Solutions to Avoid and Mitigate Environmental Impacts of Offshore Windfarms. BioConsult SH for WWF France, Husum, Germany.

Degraer, S., D. A. Carey, J. W. P. Coolen, Z. L. Hutchison, F. Kerckhof, B. Rumes, and J. Vanaverbeke. 2020. Offshore wind farm artificial reefs affect ecosystem structure and functioning: A synthesis. Oceanography 33:48–57.

Dias, M. P., R. Martin, E. J. Pearmain, I. J. Burfield, C. Small, R. A. Phillips, O. Yates, B. Lascelles, P. G. Borboroglu, and J. P. Croxall. 2019. Threats to seabirds: A global assessment. Biological Conservation 237:525–537.

Dierschke, V., R. W. Furness, and S. Garthe. 2016. Seabirds and offshore wind farms in European waters: Avoidance and attraction. Biological Conservation 202:59–68.

Dorey, K., S. Dickey, and T. R. Walker. 2019. Testing efficacy of bird deterrents at wind turbine facilities: a pilot study in Nova Scotia, Canada. Proceedings of the Nova Scotian Institute of Science (NSIS) 50:91.

Drewitt, A. L., and R. H. W. Langston. 2006. Assessing the impacts of wind farms on birds. Ibis 148:29–42.

Drewitt, A. L., and R. H. W. Langston. 2008. Collision effects of wind-power generators and other obstacles on birds. Annals of the New York Academy of Sciences 1134:233–66.

Dwyer, J. F., A. K. Pandey, L. A. McHale, and R. E. Harness. 2019. Near-ultraviolet light reduced Sandhill Crane collisions with a power line by 98%. The Condor: Ornithological Applications 121:duz008.

English, P. A., T. I. Mason, J. T. Backstrom, B. J. Tibbles, A. A. Mackay, M. J. Smith, and T. Mitchell. 2017. Improving Efficiencies of National Environmental Policy Act Documentation for Offshore Wind Facilities Case Studies Report. US Dept. of the Interior, Bureau of Ocean Energy Management, Sterling, VA.

Enos, J. K., M. P. Ward, and M. E. Hauber. 2021. A review of the scientific evidence on the impact of biologically salient frightening devices to protect crops from avian pests. Crop Protection 148:105734.

Environment and Climate Change Canada. 2016. Procedures for handling and documenting stranded birds encountered on infrastructure offshore Atlantic Canada. Environment and Climate Change Canada, Gatineau, Quebec.

Eshleman, K. N., and A. Elmore. 2013. Recommended Best Management Practices for Marcellus Shale Gas Development in Maryland. Maryland Department of the Environment, Baltimore, Maryland.

E-TWG Bird and Bat Specialist Committee. 2020. Summary of Discussions from the Bird and Bat Specialist Committee of the Environmental Technical Working Group (E-TWG). New York Offshore Wind Environmental Technical Working Group, Albany, NY.

Evans, W. R., Y. Akashi, N. S. Altman, and A. M. Manville. 2007. Response of night-migrating songbirds in cloud to colored and flashing light. North American Birds 60:476–488.

Everaert, J., and E. W. M. Stienen. 2007. Impact of wind turbines on birds in Zeebrugge (Belgium): Significant effect on breeding tern colony due to collisions. Biodiversity and Conservation 16:3345–3359.

Felton, S. K., L. P. Perkins, J. P. Smith, and S. B. Terrell. 2024. Evaluating the Effectiveness of a Detection and Deterrent System in Reducing Golden Eagle Fatalities at Operational Wind Facilities. Renewable Energy Wildlife Institute, Washington, D.C.

Ferrer, M., A. Alloing, R. Baumbush, and V. Morandini. 2022. Significant decline of Griffon Vulture collision mortality in wind farms during 13-year of a selective turbine stopping protocol. Global Ecology and Conservation 38:e02203.

Fox, A. D., and I. K. Petersen. 2019. Offshore wind farms and their effects on birds. Dansk Orn. Foren. Tidsskr. 113:86–101.

Frick, W. F., T. Kingston, and J. Flanders. 2020. A review of the major threats and challenges to global bat conservation. Annals of the New York Academy of Sciences 1469:5–25.

Garcia-Rosa, P. B., and J. O. G. Tande. 2023. Mitigation measures for preventing collision of birds with wind turbines. Journal of Physics: Conference Series 2626:012072.

Garthe, S., H. Schwemmer, V. Peschko, N. Markones, S. Müller, P. Schwemmer, and M. Mercker. 2023. Large-scale effects of offshore wind farms on seabirds of high conservation concern. Scientific Reports 13:4779.

Gartman, V., L. Bulling, M. Dahmen, G. Geißler, and J. Köppel. 2016. Mitigation measures for wildlife in wind energy development, consolidating the state of knowledge — Part 2: Operation, decommissioning. Journal of Environmental Assessment Policy and Management 18:1650014.

Garvin, J. C., J. L. Simonis, and J. L. Taylor. 2024. Does size matter? Investigation of the effect of wind turbine size on bird and bat mortality. Biological Conservation 291:110474.

Gaultier, S. P., T. M. Lilley, E. J. Vesterinen, and J. E. Brommer. 2023. The presence of wind turbines repels bats in boreal forests. Landscape and Urban Planning 231:104636.

Gauthreaux, S., and C. G. Belser. 2006. Effects of artificial night lighting on migrating birds. Pages 67–93 in. Ecological Consequences of Artificial Night Lighting. Island Press, Washington, D.C.

Gehring, J., P. Kerlinger, and A. M. Manville. 2009. Communication towers, lights, and birds: successful methods of reducing the frequency of avian collisions. Ecological Applications 19:505–514.

Gilmour, L. R. V., M. W. Holderied, S. P. C. Pickering, and G. Jones. 2020. Comparing acoustic and radar deterrence methods as mitigation measures to reduce human-bat impacts and conservation conflicts. M. Smotherman, editor. PLOS ONE 15:e0228668.

Gilmour, L. R. V., M. W. Holderied, S. P. C. Pickering, and G. Jones. 2021. Acoustic deterrents influence foraging activity, flight and echolocation behaviour of free-flying bats. The Journal of Experimental Biology 224:jeb242715.

Gilsdorf, J. M., S. E. Hygnstrom, and K. C. VerCauteren. 2002. Use of frightening devices in wildlife damage management. Integrated Pest Management 7:29–45.

Glahn, J. F., G. Ellis, P. Fioranelli, and B. S. Dorr. 2000. Evaluation of Moderate and Low-Powered Lasers for Dispersing Double-crested Cormorants from Their Night Roosts. Pages 34–45 in. Proceedings of the 9th Wildlife Damage Management Conference. Volume 11.

Goodale, M. W., and A. Milman. 2020. Assessing cumulative exposure of Northern Gannets to offshore wind farms. Wildlife Society Bulletin 44:252–259.

Goodale, M. W., and I. J. Stenhouse. 2016. A conceptual model to determine vulnerability of wildlife populations to offshore wind energy development. Human-Wildlife Interactions 10:53–61.

Gorresen, M. M., P. M. Cryan, D. C. Dalton, S. Wolf, J. A. Johnson, C. M. Todd, and F. J. Bonaccorso. 2015. Dim Ultraviolet Light as a Means of Deterring Activity by the Hawaiian Hoary Bat *Lasiurus cinereus semotus*. Endangered Species Research 28:249–257.

Gottlieb, I., T. D. Allison, C. Donovan, M. Whitby, and L. New. 2024. Developing and Evaluating a Smart Curtailment Strategy Integrated with a Wind Turbine Manufacturer Platform. Renewable Energy Wildlife Institute (REWI), Washington, DC (United States).

Guest, E. E., B. F. Stamps, N. D. Durish, A. M. Hale, C. D. Hein, B. P. Morton, S. P. Weaver, and S. R. Fritts. 2022. An Updated Review of Hypotheses Regarding Bat Attraction to Wind Turbines. Animals 12.

Hayes, M. A., L. A. Hooton, K. L. Gilland, C. Grandgent, R. L. Smith, S. R. Lindsay, J. D. Collins, S. M. Schumacher, P. A. Rabie, J. C. Gruver, and J. Goodrich-Mahoney. 2019. A smart curtailment approach for reducing bat fatalities and curtailment time at wind energy facilities. Ecological Applications 0:1–18.

Hazel, J., I. R. Lawler, H. Marsh, and S. Robson. 2007. Vessel speed increases collision risk for the green sea turtle *Chelonia mydas*. Endangered Species Research 3:105–113.

Hein, C., K. A. Williams, and E. Jenkins. 2021. Bat Workgroup Report for the State of the Science Workshop on Wildlife and Offshore Wind Energy 2020: Cumulative Impacts. New York Energy Research and Development Authority, Albany, NY.

Hill, R., K. Hill, R. Aumuller, A. Schulz, T. Dittmann, C. Kulemeyer, and T. Coppack. 2014. Of birds, blades, and barriers: Detecting and analysing mass migration events at alpha ventus. Pages 111–132 *in* A. Beiersdorf and K. Wollny-Goerke, editors. Ecological Research at the Offshore Windfarm alpha ventus - Challenges, Results, and Perspectives. Springer Spektrum, Hamburg and Berlin, Germany.

Hodos, W. 2003. Minimization of Motion Smear: Reducing Avian Collisions with Wind Turbines. National Renewable Energy Laboratory, Golden, CO.

Houlahan, J. E., S. T. McKinney, T. M. Anderson, and B. J. McGill. 2017. The priority of prediction in ecological understanding. Oikos 126:1–7.

H.T. Harvey & Associates. 2018. Evaluating a Commercial-Ready Technology for Raptor Detection and Deterrence at a Wind Energy Facility in California. American Wind Wildlife Institute, Washington, D.C.,.

Huso, M., and D. Dalthorp. 2023. Reanalysis indicates little evidence of reduction in eagle mortality rate by automated curtailment of wind turbines. Journal of Applied Ecology 60:2282–2288.

Jägerbrand, A. K., and C. A. Bouroussis. 2021. Ecological Impact of Artificial Light at Night: Effective Strategies and Measures to Deal with Protected Species and Habitats. Sustainability 13:5991.

Jain, A., R. Koford, A. Hancock, and G. Zenner. 2011. Bat mortality and activity at a northern Iowa wind resource area. The American Midland Naturalist 165:185–200.

Jodice, P. G. R., E. M. Adams, J. Lamb, Y. Satgé, and J. S. Gleason. 2019. GoMAMN Strategic Bird Monitoring Guidelines: Seabirds. Pages 129–170 in. Strategic Bird Monitoring Guidelines for the Northern Gulf of Mexico. Mississippi Agricultural and Forestry Experiment Station Research Bulletin, Mississippi State University.

Johnson, G. D., W. P. Erickson, M. D. Strickland, M. F. Shepherd, D. A. Shepherd, and S. A. Sarappo. 2003. Mortality of Bats at a Large-scale Wind Power Development at Buffalo Ridge, Minnesota. American Midland Naturalist 150:332–342.

Jonasson, K. A., A. M. Adams, A. F. Brokaw, M. D. Whitby, M. T. O’Mara, and W. F. Frick. 2024. A multisensory approach to understanding bat responses to wind energy developments. Mammal Review 54:229–242.

Jonasson, K. A., A. J. Corcoran, L. Dempsey, T. J. Weller, and J. Clerc. 2025. Bats flying through a Y-maze are visually attracted to wind turbine surfaces. Biology Letters 21:20250242.

Kerlinger, P., J. L. Gehring, W. P. Erickson, R. Curry, A. Jain, and J. Guarnaccia. 2010. Night migrant fatalities and obstruction lighting at wind turbines in North America. The Wilson Journal of Ornithology 122:744–754.

Lagerveld, S., B. J. Poerink, R. Haselager, and H. Verdaat. 2014. Bats in Dutch offshore wind farms in autumn 2012. Lutra 57:61–69.

Lagerveld, S., T. Wilkes, M. E. B. van Puijenbroek, B. C. A. Noort, and S. C. V. Geelhoed. 2023. Acoustic monitoring reveals spatiotemporal occurrence of Nathusius’ pipistrelle at the southern North Sea during autumn migration. Environmental Monitoring and Assessment 195:1016.

Lamb, J., J. Gulka, E. Adams, A. Cook, and K. A. Williams. 2024. A synthetic analysis of post-construction displacement and attraction of marine birds at offshore wind energy installations. Environmental Impact Assessment Review 108:107611.

Langston, R. H. W. 2013. Birds and wind projects across the pond: A UK perspective. Wildlife Society Bulletin 37:5–18.

Langston, R., and J. Pullan. 2003. Windfarms and Birds: An analysis of the effects of windfarms on birds, and guidance on environmental assessment criteria and site selection issues. BirdLife International to the Council of Europe, Bern Convention on the Conservation of European Wildlife and Natural Habitats, Strasbourg, Germany.

Levrel, H., S. Pioch, and R. Spieler. 2012. Compensatory mitigation in marine ecosystems: Which indicators for assessing the “no net loss” goal of ecosystem services and ecological functions? Marine Policy 36:1202–1210.

Liang, X., C. Yang, Y. Zhang, and Y. Xue. 2023. Mitigating the Negative Impact of Wind Power on Soaring Birds through Government Restrictions. Energies 16:6584.

Lüdeke, J. 2017. Offshore wind energy: Good practice in impact assessment, mitigation and compensation. Journal of Environmental Assessment Policy and Management 19:1750005.

Lüdeke, J. 2019. Retrospective and Outlook for an Environmentally Sound Development of Offshore Wind Energy. International Journal of Environmental Science 4:1–19.

Maclean, I. M. D., M. M. Rehfisch, H. Skov, and C. B. Thaxter. 2013. Evaluating the statistical power of detecting changes in the abundance of seabirds at sea. M. Bolton, editor. Ibis 155:113–126.

Marques, A. T., H. Batalha, S. Rodrigues, H. Costa, M. João, R. Pereira, C. Fonseca, M. Mascarenhas, and J. Bernardino. 2014. Understanding bird collisions at wind farms: An updated review on the causes and possible mitigation strategies. Biological Conservation 179:40–52.

Martins, R. C., J. Bernardino, and F. Moreira. 2023. A review of post-construction monitoring practices used in the evaluation of transmission power line impacts on birds and mitigation effectiveness, with proposals for guideline improvement. Environmental Impact Assessment Review 100:107068.

May, R. 2017. Mitigation for birds. Wildlife and Wind Farms - Conflicts and Solutions: Onshore: Monitoring and Mitigation. Pelagic Publishing.

May, R., T. Nygård, U. Falkdalen, J. Åström, Ø. Hamre, and B. G. Stokke. 2020. Paint it black: Efficacy of increased wind turbine rotor blade visibility to reduce avian fatalities. Ecology and Evolution 10:8927–8935.

May, R., O. Reitan, K. Bevanger, S.-H. Lorentsen, and T. Nygård. 2015. Mitigating wind-turbine induced avian mortality: Sensory, aerodynamic and cognitive constraints and options. Renewable and Sustainable Energy Reviews 42:170–181.

McClure, C. J. W., B. W. Rolek, L. Dunn, J. D. McCabe, L. Martinson, and T. Katzner. 2021. Eagle fatalities are reduced by automated curtailment of wind turbines. K. Suryawanshi, editor. Journal of Applied Ecology 58:446–452.

McClure, C. J. W., B. W. Rolek, L. Dunn, J. D. McCabe, L. Martinson, and T. E. Katzner. 2022. Confirmation that eagle fatalities can be reduced by automated curtailment of wind turbines. Ecological Solutions and Evidence 3:e12173.

McClure, C. J. W., B. W. Rolek, L. Dunn, J. D. McCabe, L. Martinson, and T. E. Katzner. 2023. Reanalysis ignores pertinent data, includes inappropriate observations, and disregards realities of applied ecology: Response to Huso and Dalthorp (2023). Journal of Applied Ecology 60:2289–2294.

McGregor, R., M. Trinder, and N. Goodship. 2022. Assessment of compensatory measures for impacts of offshore windfarms on seabirds. Natural England, York, UK.

Miles, W., S. Money, R. Luxmoore, and R. W. Furness. 2010. Effects of artificial lights and moonlight on petrels at St Kilda. Bird Study 57:244–251.

Millon, L., K. Barré, R. Julliard, P. Compère, and C. Kerbiriou. 2021. Calculation of biodiversity level between different land-uses to improve conservation outcomes of biodiversity offsetting. Land Use Policy 101:105161.

Millon, L., J.-F. Julien, R. Julliard, and C. Kerbiriou. 2015. Bat activity in intensively farmed landscapes with wind turbines and offset measures. Ecological Engineering 75:250–257.

Mockrin, M. H., and R. A. Gravenmier. 2012. Synthesis of wind energy development and potential impacts on wildlife in the Pacific Northwest, Oregon and Washington. United States Department of Agriculture, Forest Service, Pacific Northwest Research Station, Portland, OR.

Montevecchi, W. A. 2006. Influences of artificial light on marine birds. Pages 94–113 in C. Rich and T. Longcore, editors. Ecological Consequences of Artificial Night Lighting. Island Press, Washington, D.C.

Montoney, A. J., and H. C. Boggs. 1993. Effects of a Bird Hazard Reduction Force on Reducing Bird / Aircraft Strike Hazards At the Atlantic City International Airport, NJ. Pages 1–8 in. Proceedings of the 6th Eastern Wildlife Damage Control Conference.

Mott, D. F., and S. K. Timbrook. 1988. Alleviating Nuisance Canada Goose Problems with Acoustical Stimuli. Proceedings of the Thirteenth Vertebrate Pest Conference 301–304.

Murgatroyd, M., W. Bouten, and A. Amar. 2021. A predictive model for improving placement of wind turbines to minimise collision risk potential for a large soaring raptor. Journal of Applied Ecology 58:857–868.

Neate-Clegg, M. H. C., J. J. Horns, F. R. Adler, M. Ç. Kemahlı Aytekin, and Ç. H. Şekercioğlu. 2020. Monitoring the world’s bird populations with community science data. Biological Conservation 248:108653.

Nicholls, B., and P. A. Racey. 2007. Bats Avoid Radar Installations: Could Electromagnetic Fields Deter Bats From Colliding With Wind Turbines? PLoS ONE 2:e297.

Nicholls, B., and P. A. Racey. 2009. The aversive effect of electromagnetic radiation on foraging bats - A possible means of discouraging bats from approaching wind turbines. PLoS ONE 4:e6246.

Nogales, M., E. Vidal, F. M. Medina, E. Bonnaud, B. R. Tershy, K. J. Campbell, and E. S. Zavaleta. 2013. Feral Cats and Biodiversity Conservation: The Urgent Prioritization of Island Management. BioScience 63:804–810.

O’Neil, D. R. 2020. Reducing Bat Fatalities Using Ultrasonic Acoustic Deterrent Technology: A Potential Mechanism for Conservation at Offshore Wind Energy Sites. M.S. thesis, Harvard Extension School, Boston, MA.

Orloff, S., and A. Flannery. 1992. Wind Turbine Effects on Avian Activity, Habitat Use, and Mortality in Altamont Pass and Solano Count Wind Resource Areas: 1989-1991. A report prepared for California Energy Commission. BioSystems Analysis, Inc., Tiburon, CA.

Pascal, M., O. Lorvelec, V. Bretagnolle, and J. Culioli. 2008. Improving the breeding success of a colonial seabird: a cost-benefit comparison of the eradication and control of its rat predator. Endangered Species Research 4:267–276.

Peste, F., A. Paula, L. P. da Silva, J. Bernardino, P. Pereira, M. Mascarenhas, H. Costa, J. Vieira, C. Bastos, C. Fonseca, and M. J. R. Pereira. 2015. How to mitigate impacts of wind farms on bats? A review of potential conservation measures in the European context. Environmental Impact Assessment Review 51:10–22.

Peterson, T., A. Rusk, S. Aghababian, S. Edwards, and C. Byrne. 2025. Acoustic exposure reveals variation in curtailment effectiveness at reducing bat fatality at wind turbines. Ecosphere 16:e70277.

Phalan, B., G. Hayes, S. Brooks, D. Marsh, P. Howard, B. Costelloe, B. Vira, A. Kowalska, and S. Whitaker. 2018. Avoiding impacts on biodiversity through strengthening the first stage of the mitigation hierarchy. Oryx 52:316–324.

Poot, H., B. J. Ens, H. de Vries, M. A. H. Donners, M. R. Wernand, and J. M. Marquenie. 2008. Green light for nocturnally migrating birds. Ecology & Society 13:47.

Rebke, M., V. Dierschke, C. N. Weiner, R. Aumüller, K. Hill, and R. Hill. 2019. Attraction of nocturnally migrating birds to artificial light: The influence of colour, intensity and blinking mode under different cloud cover conditions. Biological Conservation 233:220–227.

Reed, J. R., J. L. Sincock, and J. P. Hailman. 1985. Light Attraction in Endangered Procellariiform Birds : Reduction by Shielding Upward Radiation. The Auk 102:377–383.

Regional Synthesis Workgroup of the Environmental Technical Working Group. 2023. Responsible Practices for Regional Wildlife Monitoring and Research in Relation to Offshore Wind Energy Development. New York Offshore Wind Environmental Technical Working Group, Albany, NY.

Reusch, C., M. Lozar, S. Kramer-Schadt, and C. C. Voigt. 2022. Coastal onshore wind turbines lead to habitat loss for bats in Northern Germany. Journal of Environmental Management 310:114715.

Ronconi, R. A., K. A. Allard, and P. D. Taylor. 2015. Bird interactions with offshore oil and gas platforms: Review of impacts and monitoring techniques. Journal of Environmental Management 147:34–45.

Ronconi, R. A., and C. C. St. Clair. 2002. Management Options to Reduce Boat Disturbance on Foraging Black Guillemots (*Cepphus grylle*) in the Bay of Fundy. Biological Conservation 108:265–271.

Russell, D. J. F., S. M. J. M. Brasseur, D. Thompson, G. D. Hastie, V. M. Janik, G. Aarts, B. T. McClintock, J. Matthiopoulos, S. E. W. Moss, and B. McConnell. 2014. Marine mammals trace anthropogenic structures at sea. Current Biology 24:638–639.

Rydell, J., H. Engström, A. Hedenström, J. K. Larsen, J. Pettersson, and M. Green. 2012. The effect of wind power on birds and bats-A synthesis. Swedish Environmental Protection Agency, Stockholm, Sweden.

Schirmacher, M. R. 2020. Evaluating the Effectiveness of an Ultrasonic Acoustic Deterrent in Reducing Bat Fatalities at Wind Energy Facilities. Bat Conservation International, Austin, TX.

Seewagen, C. L., and A. M. Adams. 2021. Turning to the dark side: LED light at night alters the activity and species composition of a foraging bat assemblage in the northeastern United States. Ecology and Evolution 11:5635–5645.

Sherman, D. E., and A. E. Barras. 2004. Efficacy of a Laser Device for Hazing Canada Geese from Urban Areas of Northeast Ohio. The Ohio Journal of Science, 104:38–42.

Simmons, R. E., S. Ralston-Paton, R. Colyn, and M.-S. Garcia-Heras. 2020. Black Harriers and Wind Energy: Guidelines for impact assessment, monitoring and mitigation. BirdLife South Africa, Johannesburg, South Africa.

Sinclair, K., and E. DeGeorge. 2016. Framework for Testing the Effectiveness of Bat and Eagle Impact-Reduction Strategies at Wind Energy Projects. National Renewable Energy Laboratory, U.S. Department of Energy, Golden, CO.

Skov, H., S. Heinanen, T. Norman, R. M. Ward, S. Mendez-Roldan, and I. Ellis. 2018. ORJIP Bird Collision and Avoidance Study. Final Report - April 2018. The Carbon Trust, United Kingdom.

Skov, H., R. S. Tjørnløv, M. Armitage, M. Barker, J. B. Jørgensen, L. O. Mortensen, and T. Uhrenholdt. 2025. High-resolution multi-sensor technology reveals low collision risk to seabirds in offshore wind farms. Journal of Applied Ecology e70239.

Smallwood, K. S., and D. A. Bell. 2020. Effects of Wind Turbine Curtailment on Bird and Bat Fatalities. Journal of Wildlife Management. 10.1002/jwmg.21844.

Smokorowski, K. E., and R. G. Randall. 2017. Cautions on using the Before-After-Control-Impact design in environmental effects monitoring programs. FACETS 2:212–232.

Solick, D. I., and C. M. Newman. 2021. Oceanic records of North American bats and implications for offshore wind energy development in the United States. Ecology and Evolution 11:14433–14447.

Spatz, D. R., N. D. Holmes, D. J. Will, S. Hein, Z. T. Carter, R. M. Fewster, B. Keitt, P. Genovesi, A. Samaniego, D. A. Croll, B. R. Tershy, and J. C. Russell. 2022. The global contribution of invasive vertebrate eradication as a key island restoration tool. Scientific Reports 12:13391.

Stantec Consulting Services Inc. 2016. Long-term bat monitoring on islands, offshore structures, and coastal sites in the Gulf of Maine, mid-Atlantic, and Great Lakes - Final Report. U.S. Department of Energy, Tompsham, ME.

Stevens, G. R., J. Rogue, R. Weber, and L. Clark. 2000. Evaluation of a Radar-activated, Demand-performance Bird Hazing System. International Biodeterioration and Biodegradation 45:129–137.

Stickley, A. R., and J. O. King. 1993. Long-Term Trial of An Inflatable Effigy Scare Device for Repelling Cormorants from Catfish Ponds. Pages 1–5 in. Proceedings of the 6th Eastern Wildlife Damage Control Conference.

Stokke, B. G., T. Nygård, U. Falkdalen, H. C. Pedersen, and R. May. 2020. Effect of tower base painting on willow ptarmigan collision rates with wind turbines. Ecology and Evolution 10:5670–5679.

Stone, E. L., S. Harris, and G. Jones. 2015. Impacts of artificial lighting on bats: A review of challenges and solutions. Mammalian Biology 80:213–219.

Sutherland, W. J., L. V. Dicks, S. O. Petrovan, and R. K. Smith. 2021. What Works in Conservation: 2021. Open Book Publishers.

Syposz, M., O. Padget, J. Willis, B. M. V. Doren, N. Gillies, A. L. Fayet, M. J. Wood, A. Alejo, and T. Guilford. 2021. Avoidance of different durations, colours and intensities of artificial light by adult seabirds. Scientific Reports 1–13.

Szostek, C. L., A. Edwards-Jones, N. J. Beaumont, and S. C. L. Watson. 2024. Primary vs grey: A critical evaluation of literature sources used to assess the impacts of offshore wind farms. Environmental Science & Policy 154:103693.

Thaxter, C. B., V. H. Ross-Smith, W. Bouten, N. A. Clark, G. J. Conway, E. A. Masden, G. D. Clewley, L. J. Barber, and N. H. K. Burton. 2019. Avian vulnerability to wind farm collision through the year: Insights from lesser black-backed gulls (*Larus fuscus*) tracked from multiple breeding colonies. Journal of Applied Ecology 56:2410–2422.

Thébaud, O., F. Boschetti, S. Jennings, A. D. M. Smith, and S. Pascoe. 2015. Of sets of offsets: Cumulative impacts and strategies for compensatory restoration. Ecological Modelling 312:114–124.

Thelander, C. G., K. S. Smallwood, and L. Rugge. 2003. Bird Risk Behaviors and Fatalities at the Altamont Pass Wind Resource Area: Period of Performance, March 1998--December 2000. National Renewable Energy Laboratory, Golden, CO.

Thompson, M., J. A. Beston, M. Etterson, J. E. Diffendorfer, and S. R. Loss. 2017. Factors associated with bat mortality at wind energy facilities in the United States. Biological Conservation 215:241–245.

Thurber, B. G., R. J. Kilpatrick, G. H. Tang, C. Wakim, and J. R. Zimmerling. 2023. Economic Impacts of Curtailing Wind Turbine Operations for the Protection of Bat Populations in Ontario. Wind 3:291–301.

Tjørnløv, R. S., H. Skov, M. Armitage, M. Barker, J. B. Jørgensen, L. O. Mortensen, K. Thomas, T. Uhrenholdt, and 11820296. 2023. Resolving Key Uncertainties of Seabird Flight and Avoidance Behaviours at Offshore Wind Farms: Final report for the study period 2020-2021. Vattbenfall, Horsholm, Denmark.

Tulp, I., H. Schekkerman, J. K. Larsen, J. Van Der Winden, R. J. W. Van De Haterd, P. Van Horssen, S. Dirksen, and A. L. Spaans. 1999. Nocturnal Flight Activity of Sea Ducks Near the Windfarm Tunø Knob in the Kattegat. Bureau Waardenburg, Culemborg, Netherlands.

U.S. Fish and Wildlife Service (USFWS). 2012. U.S. Fish and Wildlife Service Land-based Wind Energy Guidelines. U.S. Fish and Wildlife Service, Arlington, VA.

U.S. Fish and Wildlife Service (USFWS). 2021. Recommended Best Practices for Communication Tower Design, Siting, Construction, Operation, Maintenance, and Decommissioning. U.S. Fish and Wildlife Service, Migratory Bird Program, Falls Church, VA.

U.S. Fish and Wildlife Service (USFWS). 2023. U.S. Fish & Wildlife Service Mitigation Policy (Appendix 1, 501 FW 2). U.S. Fish and Wildlife Service, Arlington, VA.

Vaissière, A.-C., H. Levrel, S. Pioch, and A. Carlier. 2014. Biodiversity Offsets for Offshore Wind Farm Projects: The Current Situation in Europe. Marine Policy 48:172–183.

Verfuss, U. K., C. E. Sparling, C. Arnot, A. Judd, and M. Coyle. 2016. Review of Offshore Wind Farm Impact Monitoring and Mitigation with Regard to Marine Mammals. Pages 1175–1182 *in* A. N. Popper and A. Hawkins, editors. The Effects of Noise on Aquatic Life II. Volume 875. Advances in Experimental Medicine and Biology, Springer New York, New York, NY.

Voigt, C. C., K. Rehnig, O. Lindecke, and G. Pētersons. 2018. Migratory bats are attracted by red light but not by warm-white light: Implications for the protection of nocturnal migrants. Ecology and Evolution 1–9.

Voigt, C. C., M. Roeleke, L. Marggraf, G. Petersons, and S. L. Voigt-Heucke. 2017. Migratory bats respond to artificial green light with positive phototaxis. PLoS ONE 12:1–11.

Walsh, C., O. Hüppop, T. Karwinkel, M. Liedvogel, O. Lindecke, J. D. McLaren, H. Schmaljohann, and B. Siebenhüner. 2024.Light Pollution at Sea: Implications and Potential Hazards of Human Activity for Offshore Bird and Bat Movements in the Greater North Sea.

Watson, R. T., P. S. Kolar, M. Ferrer, T. Nygård, N. Johnston, W. G. Hunt, H. A. Smit-Robinson, C. J. Farmer, M. Huso, and T. E. Katzner. 2018. Raptor Interactions With Wind Energy: Case Studies From Around the World. Journal of Raptor Research 52:1–18.

Werber, Y., G. Hareli, O. Yinon, N. Sapir, and Y. Yovel. 2023. Drone-mounted audio-visual deterrence of bats: implications for reducing aerial wildlife mortality by wind turbines. Remote Sensing in Ecology and Conservation 9:404–419.

Whisson, D. A., and J. Y. Takekawa. 2000. Testing the Effectiveness of an Aquatic Hazing Device on Waterbirds in the San Francisco Bay Estuary of California. Waterbirds: The International Journal of Waterbird Biology 23:56–63.

Whitby, M. D., M. T. O’Mara, C. D. Hein, M. Huso, and W. F. Frick. 2024. A decade of curtailment studies demonstrates a consistent and effective strategy to reduce bat fatalities at wind turbines in North America. Ecological Solutions and Evidence 5:e12371.

Whitby, M. D., M. R. Schirmacher, and W. F. Frick. 2021. The State of the Science on Operational Minimization to Reduce Bat Fatality at Wind Energy Facilities. A report submitted to the National Renewable Energy Laboratory. Bat Conservation International, Austin, Texas.

Wiese, F. K., W. A. Montevecchi, G. K. Davoren, F. Huettmann, A. W. Diamond, and J. Linke. 2001. Seabirds at risk around offshore oil platforms in the Northwest Atlantic. Marine Pollution Bulletin 42:1285–1290.

Williams, K. A., J. Gulka, A. S. C. P. Cook, R. H. Diehl, A. Farnsworth, H. Goyert, C. Hein, P. Loring, D. Mizrahi, I. K. Petersen, T. Peterson, K. M. Press, and I. J. Stenhouse. 2024. A framework for studying the effects of offshore wind energy development on birds and bats in the Eastern United States. Frontiers in Marine Science 11:1274052.

Williams, K., E. Connelly, S. Johnson, and I. Stenhouse. 2015. Wildlife Densities and Habitat Use Across Temporal and Spatial Scales on the Mid-Atlantic Outer Continental Shelf: Final Report to the Department of Energy EERE Wind & Water Power Technologies Office, Award Number: DE-EE0005362. Biodiversity Research Institute, Portland, ME.

Winiarski, K. J., D. L. Miller, P. W. C. Paton, and S. R. McWilliams. 2014. A spatial conservation prioritization approach for protecting marine birds given proposed offshore wind energy development. Biological Conservation 169:79–88.

Wolf, S., B. Keitt, A. Aguirre-Muñoz, B. Tershy, E. Palacios, and D. Croll. 2006. Transboundary seabird conservation in an important North American marine ecoregion. Environmental Conservation 33:294–305.

World Bank Group. 2015. Environmental, Health, and Safety Guidelines for Wind Energy. The World Bank Group.

Yates, K. L., P. J. Bouchet, M. J. Caley, K. Mengersen, C. F. Randin, S. Parnell, A. H. Fielding, A. J. Bamford, S. Ban, A. M. Barbosa, C. F. Dormann, J. Elith, C. B. Embling, G. N. Ervin, R. Fisher, S. Gould, R. F. Graf, E. J. Gregr, P. N. Halpin, R. K. Heikkinen, S. Heinänen, A. R. Jones, P. K. Krishnakumar, V. Lauria, H. Lozano-Montes, L. Mannocci, C. Mellin, M. B. Mesgaran, E. Moreno-Amat, S. Mormede, E. Novaczek, S. Oppel, G. Ortuño Crespo, A. T. Peterson, G. Rapacciuolo, J. J. Roberts, R. E. Ross, K. L. Scales, D. Schoeman, P. Snelgrove, G. Sundblad, W. Thuiller, L. G. Torres, H. Verbruggen, L. Wang, S. Wenger, M. J. Whittingham, Y. Zharikov, D. Zurell, and A. M. M. Sequeira. 2018. Outstanding Challenges in the Transferability of Ecological Models. Trends in Ecology & Evolution 33:790–802.

